# Humanized Anti-PD-1 Antibodies Generated Using The Conditional Kernel-Elastic Autoencoder

**DOI:** 10.64898/2026.08.03.742579

**Authors:** Yuanjun Shi, Haote Li, Pulan Liu, Christopher G. Bunick, Shaogeng Tang, Jimin Wang, Victor S. Batista

## Abstract

The human immune system excels at generating highly effective antibodies through natural selection and somatic hypermutation, but adapting these antibodies for therapeutic use, referred to as "antibody medicine-likeness", requires careful consideration of biochemical and physiological properties. Traditional redesign methods are often slow and limited in scope. In this study, we introduce a machine learning-based approach to evolve new anti-PD-1 antibodies within a chemically informed latent space using a conditional kernel-elastic autoencoder (CKEA) between nivolumab and pembrolizumab, both of which bind the FG-loop "hotspot" of PD-1 in the most distantly related orientations, differing by 174°. This generative framework is designed to preserve favorable therapeutic features while exploring variants with different potency, ultimately for improved potency. To evaluate structural and functional viability, we performed molecular dynamics (MD) simulations of the generated antibody – PD-1 complexes and described their MD properties. These simulations reveal detailed free-energy landscapes and identify stable binding conformations, providing a strong basis for experimental validation. To validate our designs, we expressed and experimentally tested the antibodies for binding affinity to PD-1. Upon expression and purification, three out of six designed antibodies exhibited some binding to PD-1, whose properties could likely be improved using other computational saturation mutagenesis or laboratory evolution. Our results demonstrate the potential of artificial intelligence (AI)-guided interpolation methods to generate novel, high-affinity antibodies with therapeutic promise, offering a powerful strategy for next-generation antibody development.

**SYNOPSIS TOC:** Combination of MD simulations with machine learning algorithms could revolutionize the antibody-breeding sciences to lead to new antibody discovery that is compatible with or better than naturally occurring antibodies.

**Topics of Content (TOC):** Fingerprints of R86 finger of PD-1 are recognized by our designed P2N_2 anti-PD-1 antibody according to our MD simulations

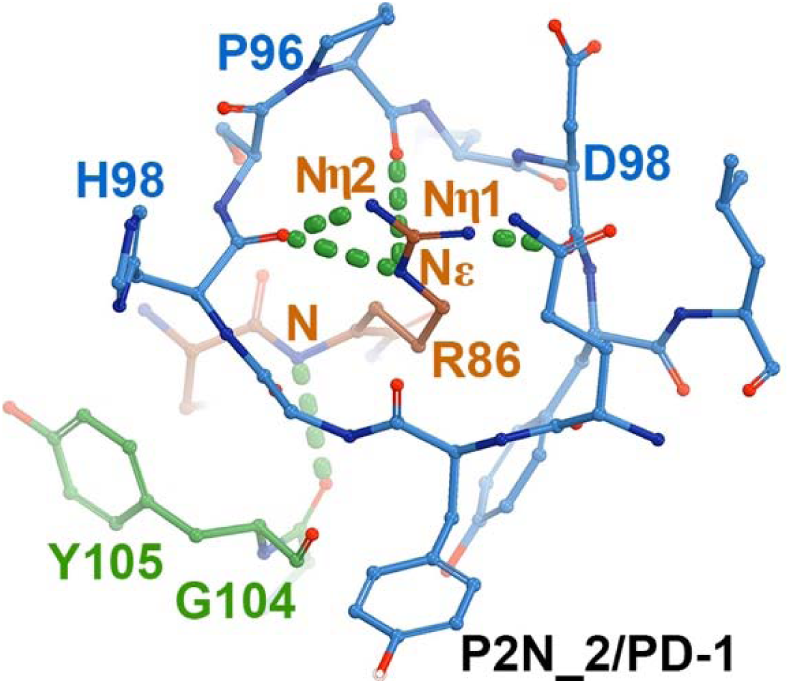

## INTRODUCTION

Programmed cell death protein 1 (PD-1) plays a critical role in immune regulation and is essential for survival in mouse models infected with various viruses, underscoring its importance in the immune response.^1, 2^ Its ligand, PD-ligand-1 (PD-L1), is frequently overexpressed in a wide range of cancers, allowing tumor cells to evade immune surveillance by binding to PD-1 on T cells and forming a suppressive PD-L1/PD-1 complex.^3–7^ This mechanism is also present in certain healthy tissues, where it helps prevent unintended immune activation, highlighting the delicate balance between immune response and immune tolerance maintained through PD-L1/PD-1 interactions. Disruption of this complex can restore T cell activity against tumor cells, making PD-1 and PD-L1 pivotal targets in the development of antibody-based cancer immunotherapies.^3–6, 8, 9^ The class of the anti-PD-1 antibodies that block PD-L1 binding are also known as antagonistic antibodies.

In this study, we aim to enhance the effectiveness of nivolumab (BMS-936558 or ONO-4538), a therapeutic antibody targeting PD-1 with binding affinity of approximately 4 nM, As a reference, we use pembrolizumab (MK-3475, formerly known as lambrolizumab), which exhibits significantly higher affinity at 27 pM (Fig. 1).^10–14^ Our objective is to design antibodies with improved binding affinity compared to nivolumab, thereby enabling lower dosing while minimizing the risk of overactivation that could adversely affect non-target cells.

**Figure 1.**
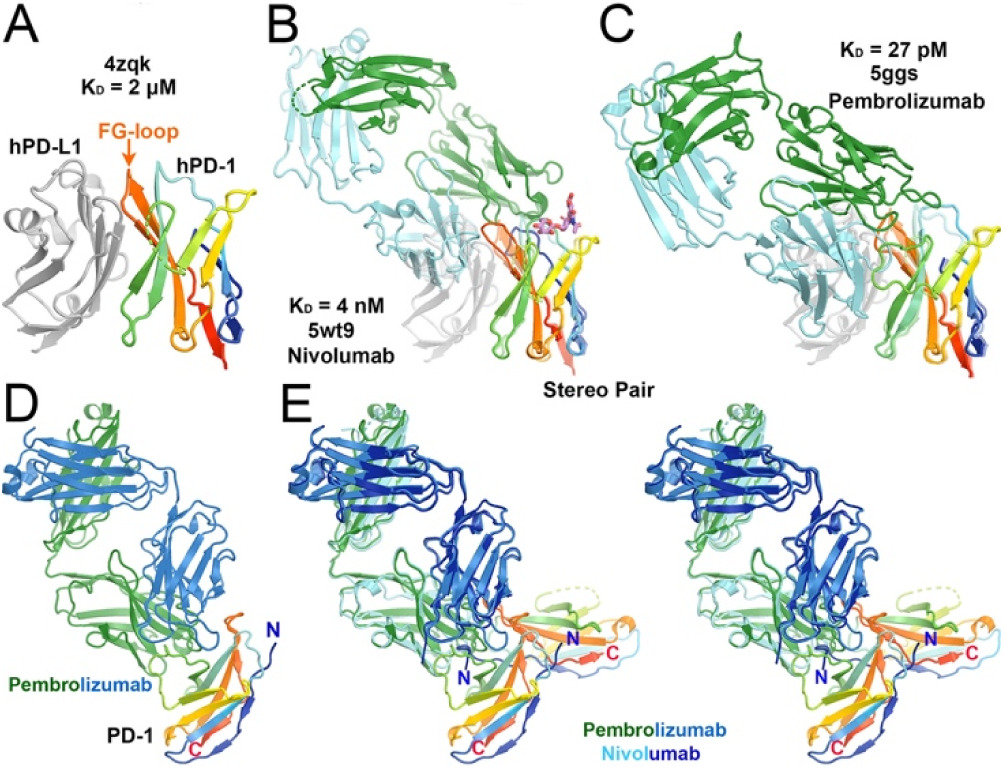
Crystal structures of human PD-1 complexes with PD-L1, or nivolumab, or pembrolizumab. (A) PD-1 (rainbow colors)/PD-L1(silver). (B) Superposition of the PD-1/PD-L1 complex with nivolumab (heavy chain in green and light chain in sky blue). (C) Superimposition of the PD-1/PD-L1 complex with pembrolizumab (green/sky blue). (D) The pembrolizumab/PD-1 complex by itself with N and C termini of PD-1 labeled. (E) Stereodiagram of antibody aligned comparison between pembrolizumab and nivolumab showing that two PD-1 (in rainbow colors) proteins have rotated by 173.6°. Binding affinity and corresponding PDB IDs are indicated. PD-1 in all three panels is oriented the same way.

Therapeutic antibodies used in long-term cancer treatment (up to 24 months) differ significantly from those employed for short-term treatment of bacterial or viral infections, or those generated in transgenic mice in response to single-dose antigen. A key factor influencing therapeutic efficacy is the *in vivo* half-life of antibodies; longer half-lives contribute positively to what is often referred to as "antibody medicine-likeness."

Pembrolizumab was discovered using transgenic mice immunized with a human PD-1/human Fc fusion followed by structure-guided redesign to introduce the S228P mutation, which mitigated unwanted reactivity to the Fc fragment used in the antigen.^10^ This antibody exhibits high specificity for human PD-1 and does not cross-react with PD-1 homologs such as CD28, CTKL-4 and ICOS (inducible T-cell Co-Stimulator), or with murine PD-1. However, most naturally occurring antibodies, including both nivolumab and pembrolizumab, many are typically short-lived when produced in response to single-shot antigens, relying on immune memory to produce effective antibodies when reencountering the same antigens in future. They are rapidly generated but also rapidly degraded upon complexation with their antigen for neutralization. As such, reengineering antibodies to enhance both half-life and higher binding affinity remains a key focus in therapeutic antibody development.^15, 16^ In this work, we explore AI-driven methods aimed at improving some of those properties, using PD-1 targeting antibodies as a representative application.

Our strategy for generating new antibodies focuses on improving nivolumab by sampling candidates through interpolation between nivolumab and pembrolizumab within the latent space of the conditional kernel-elastic autoencoder (CKEA).^17, 18^ While demonstrated using this specific antigen-antibody pair in this study, the approach can be applied using other reference points in latent space and can be extended for applications to other targets.^19–21^ The CKEA model was trained using the Feldhaus library that contains 130,000 antibody sequences,^22^ enabling the generation of humanized antibodies with properties consistent with naturally occurring sequences. To evaluate the generated candidates, we apply a set of six criteria commonly associated with therapeutic antibodies, collectively referred to as "antibody medicine- likeness".^22^

We leverage the overlapping binding sites of nivolumab and pembrolizumab on PD-1, along with their geometric relationship to PD-L1, to generate homologous models for new antibody designs. This strategy circumvents the challenges associated with *de novo* modeling, which is often prone to inaccuracies. Using MD simulations, we anticipate that the newly designed antibodies will closely resemble their respective templates, nivolumab or pembrolizumab (which contain hidden information about PD-1 hotspots), in both structure and function, supporting the feasibility of this interpolation-based approach to antibody design.

Our approach based on the CKEA differs from other methods applied to antibody design, including other types of machine learning algorithms,^23–25^ and is complemented with experimental validation. To validate our designs, we expressed and experimentally tested the antibodies for binding affinity to PD-1.^23, 26, 27^ Upon expression and purification, three out of six designed antibodies exhibited some binding to PD-1. Combination of this finding with our recent advancement on computational saturation mutagenesis, it should be possible to improve binding properties of these low-affinity designed antibodies through both computational and experimental affinity maturation procedures.^28–30^

In the past few years, a new class of anti-PD-L1 antibodies were discovered, known as agonistic antibodies.^31–33^ They do not necessarily block interactions between PD-1 and PD-L1. Instead, they interfere with interactions of PD-1 with T-cell receptors (TCRs), which initiates immune receptor signaling and therefore they are potential treatments for autoimmunity. Any computational discovery of novel anti-PD-1 antibodies could be tailored for either anti-cancer or anti-autoimmunity treatment when their specific functions can be properly defined.

Our approach also differs from many other deep-learning algorithms recently developed within the past decade or so.^34–49^ Some of these algorithms can be combined with (or integrated into) ours. Other algorithms often emphasize generalized protein-protein interactions and/or protein-drug interactions through electrostatic and shape complementarity, which are gradually extended to apply antibody-antigen interactions, a specialized type of protein-protein interactions involving very flexible surface or unstructured surfaces when both proteins are not in complex with each other. Because of flexible antigen-antibody interacting surfaces, the resolution of antibody-antigen crystal structures is often modest and the corresponding complex structures are not accurate enough for antibody redesign, a serious problem that has been overlooked by the scientific community. For example, we have succeeded in redesign of pembrolizumab with a computational study of affinity maturation only when starting with the much more reliable MD simulations-derived complex structure but we failed when directly starting with the published complex crystal structure.^28^ The detailed analysis of the MD-derived pembrolizumab/PD-1 and nivolumab/PD-1 complex structures used in the previous study,^28^ and their comparison with corresponding crystal structures are provided below in this study.

## METHODS AND MATERIALS

### Sequence data preparation and model training

Variable heavy (VH) and variable light (VL) chain regions from approximately 130,000 antibodies were extracted from a subset of the Feldhaus antibody library.^22^ In the FASTA format, each sequence was a concatenation of VL and VH sequences with a special separator token formatted as [VL]>[VH] for input into the sequence-to-sequence CKEA model. Each input– output pair corresponded to a single antibody, resulting in one sequence entry per antibody in the dataset.

Training of the CKEA model was conducted on four NVIDIA A100 GPUs for 200,000 steps over 100 epochs. Calculated medicine-likeness scores were provided to both the encoder and decoder to condition the generation of molecules. CKEA hyperparameters λ and δ were both set to 1.^17^ Each amino acid was represented using a 256-dimensional embedding vector, and the latent space was set to 2560 dimensions. The model was optimized using the ADAM optimizer with a learning rate of 0.0002.^50^ During training, gradients were clipped to a norm of 1.0. A dropout rate of 0.1 was applied to all attention operations and embedding layers.

### Gonnet matrix for sequence alignment and mutation

The default Gonnet matrix from BioPython was used to assess sequence differences between the redesigned antibodies to the reference sequences (Table S1).^51, 52^ Gap opening and extension penalties were set to -10 and -1, respectively. The substitution scores quantify the degree of divergence of the designed anti-PD-1 antibodies from the two parental antibodies. The Gonnet matrix was scaled at 250 PAM (point-accepted mutations per 100 amino acids) units, providing a measure of the chemical distance between sequences.

### Structural modeling of designed antibodies

Six new antibodies (P2N_1 through P2N_6) were generated by linear interpolation between nivolumab and pembrolizumab in latent chemical space (Table S2). All antibody structures were predicted using AlphaFold2.^53^ To assess binding stability, two docking poses were evaluated for each antibody–PD-1 complex using molecular dynamics (MD) simulations: one based on the pembrolizumab binding mode (PDB ID: 5GGS) and the other on the nivolumab binding mode (PDB ID: 5WT9),^54, 55^ which represent the most distantly related orientation, differing by 174°. We expect that MD simulations resulted in equilibrium structures with reasonable stablility when starting from configurations of these two antibodies. For comparison, the reference crystal structure of the PD-1/PD-L1 complex (PDB ID: 4ZQK) was also included.^56^

### Molecular dynamics simulations and trajectory analysis

Docked PD-L1/designed antibody complex structures were prepared using Maestro from Schrödinger, Inc, with terminal residue capped and missing hydrogen atoms added.^57^ Each complex was solvated in a rectangular box of explicit water molecules, extending 15 Å beyond the solute in all directions. Counterions (Na^+^ or/and Cl^-^) were added to neutralize the system. Parameter and topology files were generated using the *tLEaP* module from the Amber suite.^58, 59^ MD simulations were performed using NAMD.^60^ Systems were equilibrated at 310 K (i.e., 37°C, physiologically relevant body temperature) through a three-step protocol: (i) energy minimization of the solvent only, (ii) minimization of the solvent and protein side chains, and (iii) full-system minimization. For the production runs, hydrogen mass repartitioning (HMR)^61^ was applied, enabling a 4-fs integration time-step. Each system was simulated for 400 ns.

MD trajectories were aligned using either the full complexes, antibody-only, or PD-1 using the CCP4 suite and VMD.^62, 63^ Each trajectory frame was converted to an electron density (ED) (or electrostatic potential, ESP) map, and maps across all frames were summed using CCP4, following established protocols.^28, 62, 64–67^ The resulting maps were visualized with Coot and Chimera, and equilibrium structures were fitted and refined against the MD-derived maps using Coot and Phenix.^68, 69^ Figures were generated in PyMol.^70^ Residue numbering corresponds to the 5GGS (PDB ID) complex structure whenever possible, including all insertions and deletions. Glycans were omit in MD simulations because they were not used in initial docking and because they appeared to be far away from the docked sites (Fig. 2). However, the potential influence of glycan omission on complex stability and binding predictions remains uncertain and should be investigated but it is outside the scope of this study.

**Figure 2.**
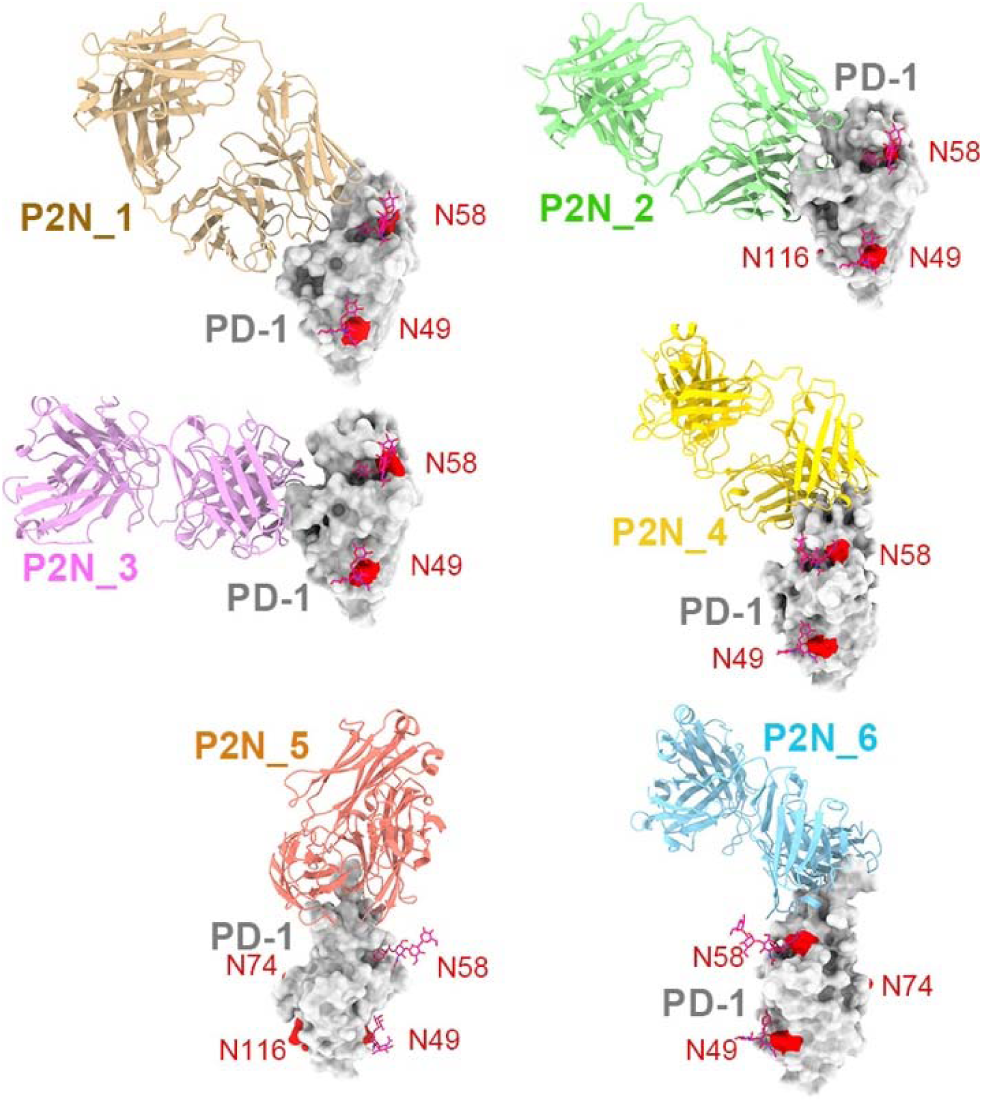
An overview of the MD-derived antibody (cartoon representation)/PD-1 (grey surface representation) complexes for six designed antibodies with known glycan shown in red balls-and-sticks.

### Molecular cloning, protein expression, purification

Oligonucleotides encoding the variable regions of designed IgG antibodies were synthesized by Integrated DNA Technologies (IDT). Designed antibodies, as well as pembrolizumab, nivolumab IgG, and Fab constructs, were cloned into the pVRC plasmid (Addgene). Constructs encoding PD-1 (C-terminally tagged with an Fc domain and AviTag), PD-L1, and PD-L2 (each with a C-terminal 6×His tag) were cloned into the pADD2 plasmid using the In-fusion snap assembly master mix (Takara).

For antibody expression, equimolar amounts of heavy and light chain plasmids were co-transfected into Expi293Fcells using FectoPro transfection reagent (Polyplus). After 5–7 days of culture, cell supernatants were harvested by centrifugation at 15,000 × g for 30 minutes.

His-tagged PD-L1, PD-L2, and Fab fragments were purified using Ni-NTA affinity chromatography and eluted with 250 mM imidazole. Eluted proteins were buffer-exchanged into storage buffer (150 mM NaCl, 20 mM HEPES, pH 7.4). AviTag-labeled IgGs and PD-1-Fc fusion proteins were purified using a 5 mL HiTrap MabSelect PrismA protein A column (Cytiva), equilibrated with the same storage buffer, and eluted with 100 mM glycine (pH 2.8). Eluates were immediately neutralized with 1 M HEPES to pH 7.4. Final protein concentrations were determined by absorbance at 280 nm using theoretical extinction coefficients calculated via the ExPASy ProtParam tool.

### Binding studies using Biolayer Interferometry (BLI) assays

All binding studies were performed using biolayer interferometry (BLI) on an Octet R8 system (Sartorius) with Octet Discovery Software 13.0 at 25Q°C (i.e., room temperature). Streptavidin (SA) biosensor tips (Sartorius) were used with a running buffer composed of 20 mM HEPES (pH 7.4), 150 mM NaCl, 0.1% BSA, and 0.05% Tween-20 (HBS-EBT). Biotinylated PD-1-Fc was immobilized onto SA biosensors to a 2 nm loading threshold. Saturation of immobilized PD-1-Fc was achieved by incubating with excess pembrolizumab IgG, nivolumab IgG, or buffer alone (control).

Association and dissociation phases were monitored for 2 minutes each in HBS-EBT buffer. After each binding cycle, biosensors were regenerated using 0.1 M glycine (pH 1.5). All experiments were performed in triplicate. Fractional binding reductions were calculated relative to the PD-1-Fc-only control using Octet Analysis Studio 13.0.

Initial kinetic parameters were derived assuming 1:1 binding and single-exponential kinetics using the built-in ForteBioanalysis software. For cases involving biphasic or multiphasic binding, data were reanalyzed using two-exponential fitting based on the method described by Kayastha and Gupta.^60^ Interpretation of fast pseudo–first-order reactions followed the frameworks of Pollard and Du.^71, 72^ The highest binding affinity (*K*_D_=*k*_off_/*k*_on_) will be the ratio between the slow-phase *k*_off_ and the fast-phase *k*_on_. The most probable *K*_D_ values were determined from the ratio between the forward rate in dominant kinetic phase and the reverse rate of the slow-phase of biphasic scheme.

## RESULTS AND DISCUSSION

### Six interpolated anti-PD-1 antibodies in latent chemical space

Both pembrolizumab and nivolumab exhibit moderate medicine-likeness scores of -1.52 and -5.52, respectively, on a scale ranging from -15 to +13 (Fig. 3).^22^ These intermediate scores suggest the presence of suboptimal physicochemical or pharmacokinetic properties (such as susceptibility to self-association, a tendency toward aggregation, or poor solubility, — factors that may reduce antibody stability and efficacy *in vivo*). While random mutagenesis is unlikely to improve these properties, our CKEA-based generative modeling approach offers a more strategic approach.

**Figure 3.**
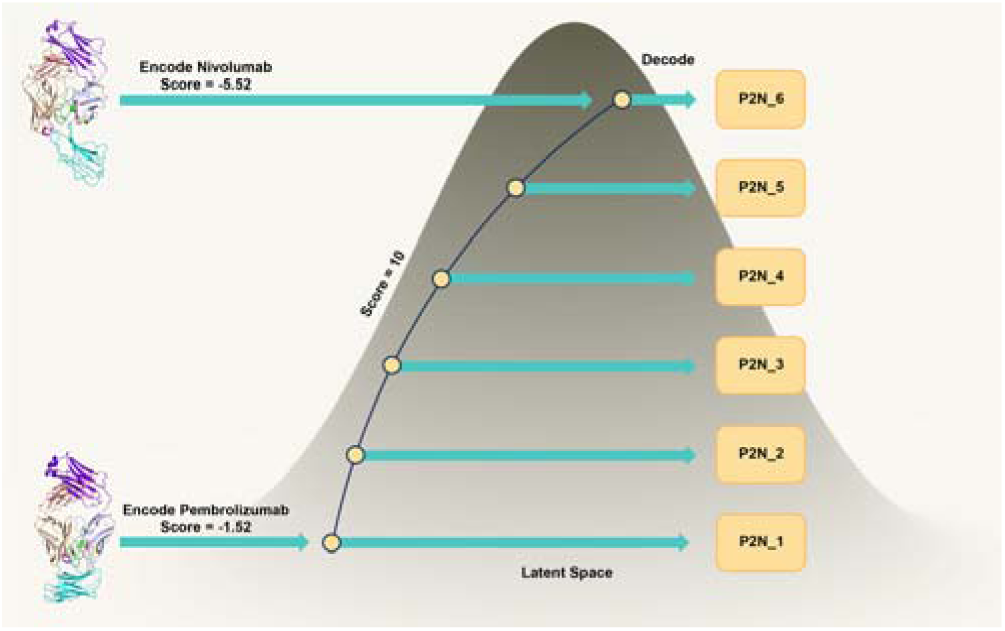
Generative antibodies with the conditioned medicine-likeness score being set 10 for interpolation between pembrolizumab (score of -1.52) and nivolumab (score of -5.52) in Gaussian-like schematic representation. Six antibodies were generated via linear interpolation in the latent chemical space.

Specifically, we conditioned our generative process (i.e., hidden information about PD-1 hotspots) to produce antibody variants, targeting a medicine-likeness score of 10 in latent space. Among the generated antibodies, P2N_1 was closest to pembrolizumab and farthest from nivolumab in latent space, whereas P2N_6 is the closet to nivolumab and the least similar to pembrolizumab (Fig. 1). The sequences of these designed antibodies, along with pairwise substitution scores relative to pembrolizumab and nivolumab (using the Gonnet substitution matrix), are provided in the Supporting Information (Tables S1–S3), with detailed comparisons within the hypervariable regions.

### MD simulations of the PD-1/PD-L1 Complex

Both pembrolizumab and nivolumab outcompete PD-L1 for binding to PD-1, with partial overlap of their binding sites although with different orientations (Fig. 1, 4). A shared feature among all known anti-PD-1 therapeutic antibodies is their targeting of the FG loop, which connects strands βF and βG of human PD-1 and is recognized as a key epitope or “hotspot” for binding of many known monoclonal antibodies.^73^ Conformation of the loop varies from one antibody complex to the next. The "hotspot" is also known to immunodominant "public epitopes" when it occurs in infectious bacteria and viruses to which germline-encoded amino acid-binding (GRAB) motifs are well represented in the database of human antibodies.^74^ Our AI-based generative approach has already taken into this hidden sequence information into account after it has been trained with the Feldhaus antibody sequence database.^22^

In addition to the FG loop, pembrolizumab also engages the C’D loop, while nivolumab uniquely interacts with the N-terminal segment of PD-1 (Fig. 4). These additional contacts lie on opposite sides of the FG loop, effectively anchoring PD-1 in two distinct orientations relative to the antibodies.

**Figure 4.**
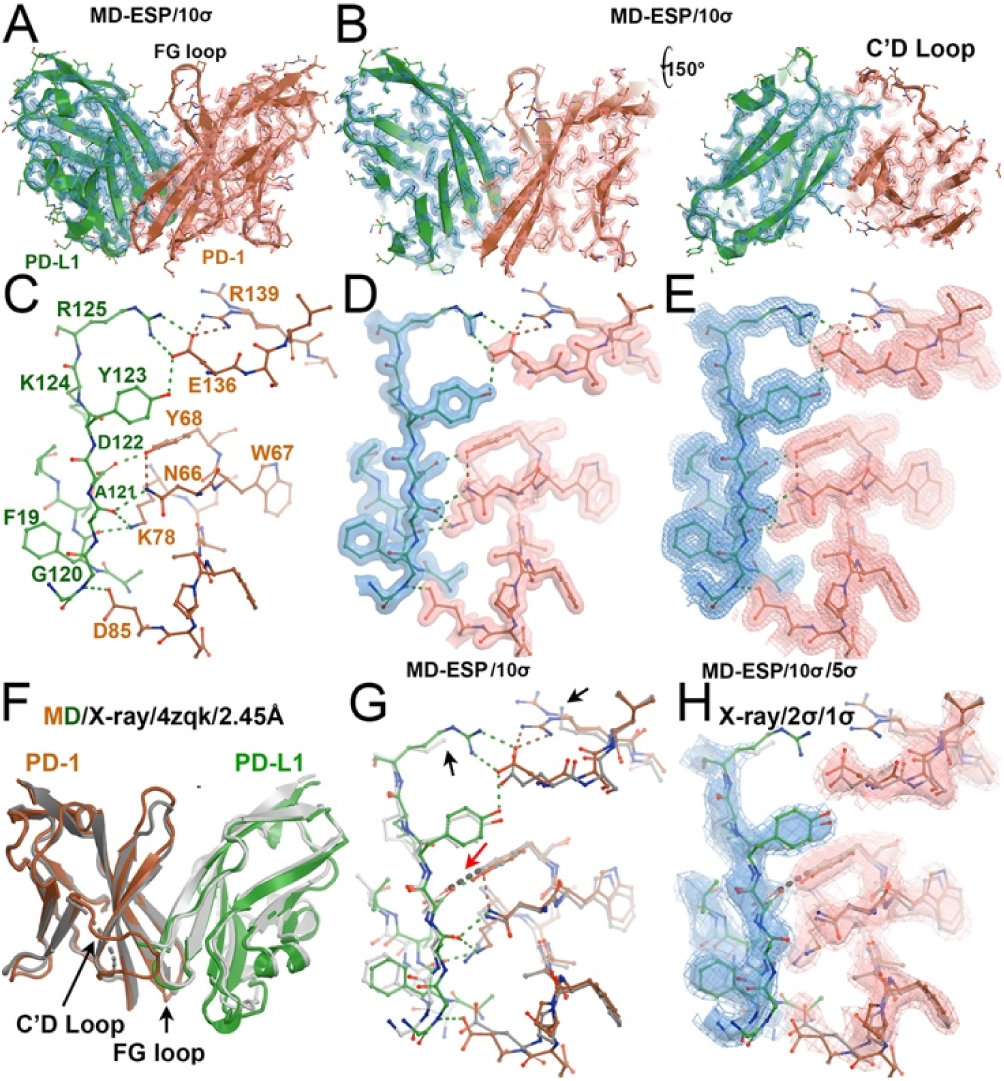
MD-derived ED maps and MD-derived structure in equilibrium of the human PD-1/PD-L1 complex and their comparison with X-ray derived ED maps and crystal structure. (A) MD-derived structure of the PD-1 (brown)/PD-L1 (green) complex superimposed onto MD-derived ED maps contoured at 10σ. (B) Front and back views in a thin slice. (C) Selected hydrogen bonds at the PD-1/PD-L1 interface. (D) With MD-ED maps contoured at 10σ. (E) With MD-ED maps contoured at both 10σ and 5σ. (F) Comparison between the MD-derived structure (brown and green) with X-ray derived crystal structure (grey and silver) at 2.45 Å resolution. Note that C’D loop was missing in the crystal structure (4zqk) and was interpretable in MD-derived maps. (G) Comparison at the subunit interface. Black arrows indicate incomplete sidechains. Red arrows show bad stereochemical clashes. (H) With X-ray derived ED maps contoured at 2σ and 1σ.

Structurally, the binding orientation of PD-1 in complexes with pembrolizumab and nivolumab differ by a 173.6° rotation about the antibody dyad axis centered on the FG loop. This makes them the most structurally divergent PD-1-antibody interactions among characterized anti-PD-1 antibodies - a key rationale for selecting these two antibodies as endpoints for linear interpolation in our design space. We anticipate that newly designed antibodies will bind somewhere between these two points.

The FG loop of PD-1 corresponds to CDR3 in the classical nomenclature of the immunoglobulin fold. In antibodies, the CDR3 regions of both heavy and light chains are located near the pseudo-dyad axis at the antigen-binding interface. The heavy chain typically features a longer CDR3, while the light chain has a shorter counterpart-hence the basis for their designation as "heavy" and "light."

Given the 173.6° rotational difference between the PD-1 binding orientations in the pembrolizumab and nivolumab complexes, initiating MD simulations from one pose (e.g., pembrolizumab-bound) and expecting convergence to the alternate pose (e.g., nivolumab-bound) is unrealistic within the 400 ns simulation timescale. Structural convergence across such a large conformational gap is kinetically limited and likely inaccessible in unbiased MD. Therefore, to ensure adequate conformational sampling, we initialized MD simulations for each of the six designed anti–PD-1 antibodies from both pembrolizumab-like and nivolumab-like binding poses. This dual-starting-point strategy increases the likelihood of capturing relevant binding conformations and reduces bias from starting geometry. It should be noted, however, that our approach cannot exclude the possibility that our designed antibodies may bind elsewhere on PD-1 that is not sampled in our analysis. Our identified binding pose in our study may not represent the highest-affinity pose.

Given that many PD-1 crystal structures were determined using *E. coli* expressed human PD-1, in which N58 remains non-glycosylated, we removed glycan moieties from crystal structures prior to conducting MD simulations. For consistency, comparisons of MD-derived ED maps were performed using non-glycosylated PD-1 whenever possible. While the role of glycosylation in antibody binding remains debated in the literature,^55, 75–78^ we suggest that some of these discrepancies may be reconciled by considering temperature-dependent binding affinities. Our MD simulations indicate that the entropic contribution (−TΔS) is often substantial in antibody-antigen interactions, implying that glycan effects could be more pronounced under specific thermodynamic conditions that may not be captured in static structures in binding assays at reduced temperature of 25°C.

MD simulations of the PD-1/PD-L1 complex, initiated from its crystal structure (PDB ID: 4ZQK), indicate that the crystallographic conformation closely resembles the structure observed under simulated physiological conditions (Fig. 4). However, the MD-derived structure reveals a more extensive and complete set of interactions between PD-1 and PD-L1, including a greater number of hydrogen-bonds and hydrophobic contacts. This enhanced interface is partly due to the inclusion of all the sidechains that are missing in the crystal structure as a result of limited resolution. For instance, in the MD-derived structure, PD-1 residue E136 forms interactions with PD-L1 R125, as well as with one of the two conformations of PD-1 R139, both of which lack defined side chains in the 4ZQK structure. Additionally, the hydrogen bond observed between Y68 of PD-1 and D122 of PD-L1 in the crystal structure features a steric clash between their side chains, which is fully resolved in the MD model. Taken together, these observations suggest that the MD-derived structure offers a more complete and stereochemical accurate representation of the PD-1/PD-L1 interaction than the reported crystal structure.

An overall distribution of MD-derived ED maps for the PD-1/PD-L1 complex closely resembles that of X-ray crystallographic ED maps, with well-defined density in the core of the complex and increased flexibility at the periphery. Notably, the FG loop of PD-1 exhibits significant mobility, as reflected by weaker density in both MD and experimental ED maps. This is consistent with its limited interactions with PD-L1 and its recognized role as a major epitope targeted by therapeutic antibodies. In contrast, the C’D loop, which connects βC’ and βD strands, is clearly resolved in the MD-derived ED maps, whereas it is completely absent in the crystal structure due to structural disorder. These differences underscore the advantage of MD simulations in capturing dynamic and flexible regions that are poorly resolved by crystallography.

Overall, the MD-derived ED map provides a more complete and stereochemical accurate representation of the PD-1/PD-L1 interface at equilibrium. The structure obtained from MD is more complete and arguably more biologically relevant than the static crystal structure. These results highlight the value of MD simulations not only as a tool for validating and interpreting experimental X-ray and cryo-EM data but also as an approach for refining structural models and uncovering otherwise inaccessible features.^79^

The MD-derived ED features for the hydrogen-bonding interaction between D122 of PD-L1 and Y68 of PD-1 are well defined and sharply localized deep within the subunit interface (Fig. 4). This suggests that deviations from the observed hydrogen bond geometry would incur a significant free energy penalty due to a highly stabilizing and structurally constrained interaction. In contrast, the hydrogen bonds formed between R125 of PD-L1 and E136 of PD-1, positioned at the periphery of the interface — are associated with less-defined ED features (Fig. 4). This suggests that the R125/E136 interaction contributes less to the overall stabilization of the complex. One likely explanation is that both R125 and E136 retain the ability to engage in favorable, solvent-mediated hydrogen-bonding, reducing their dependency on direct inter-subunit contacts. These observations support the idea that buried hydrogen-bonds, such as D122–Y68, are more structurally critical than peripheral ones like R125–E136.

### MD simulations of the PD-1/pembrolizumab complex

The crystal structure of the PD-1/pembrolizumab complex, resolved at 2.0 Å, reveals two slightly different complexes forming an antiparallel heterodimer within the crystalline lattice, which offers higher resolution and accuracy than the previously reported PD-1/PD-L1 complex at 2.45 Å resolution.^54, 56^ The antiparallel dimer arrangement observed in the crystal is likely a result of the high protein concentration used during crystallization. However, MD simulations initiated from one of the crystallographically observed complexes indicate significant conformational deviation near residue K131 of PD-1 (Fig. 5), suggesting that the crystal structure does not correspond to the configuration under physiological (lower concentration) conditions, where asymmetric dimerization is unlikely. Furthermore, PD-1 is membrane-bound and oriented parallel to the cell surface in the context of its full-length structure, including additional domains. The antiparallel dimer seen in the crystal is therefore unlikely to form *in vivo* and should be considered a crystallization artifact. It should be noted that our MD simulations were carried out at physiologically relevant temperature of 37°C while crystals were often prepared 25°C or lower (some at 4°C) and the crystal structures were determined at liquid nitrogen temperature.

**Figure 5.**
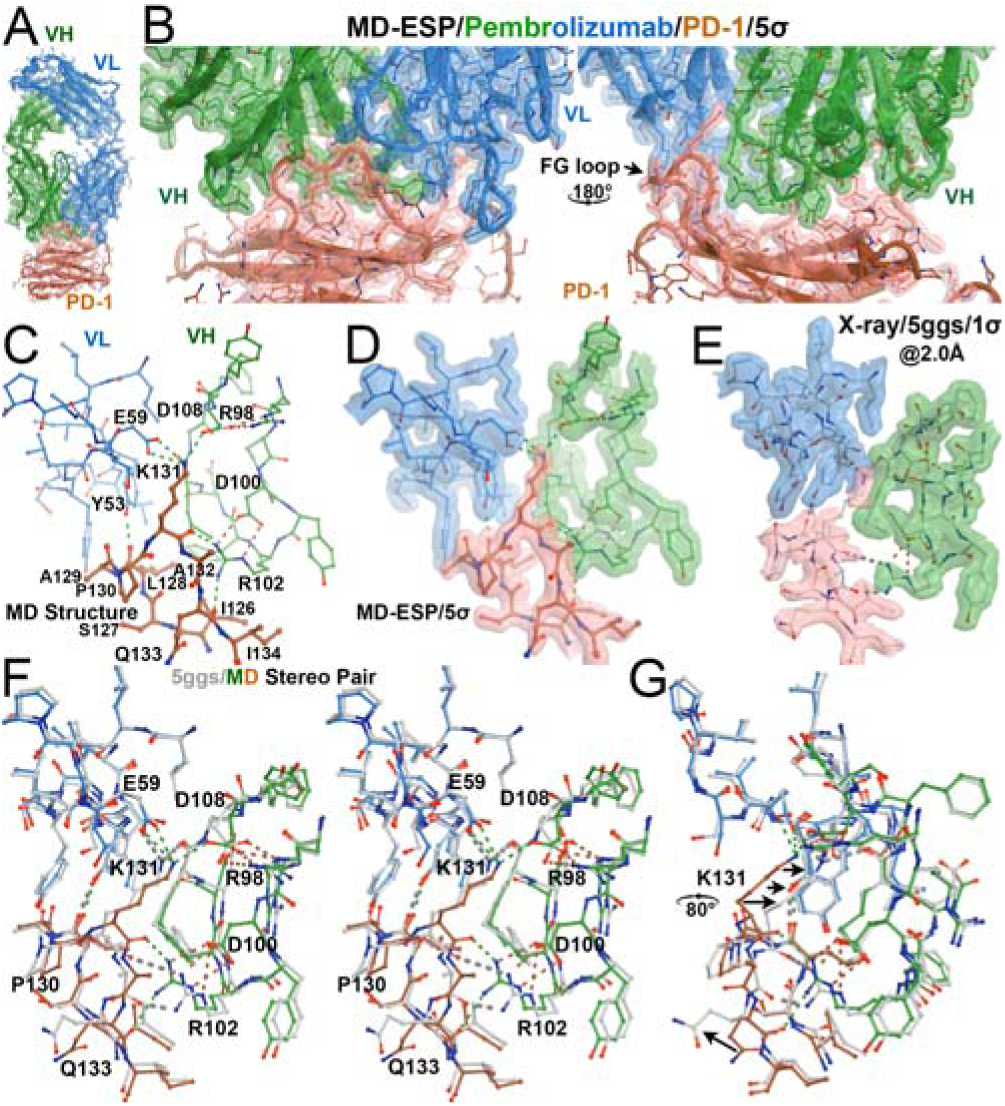
MD-derived structure of the PD-1/pembrolizumab complex and comparison with the crystal structure. (A) An overview with MD-derived ED maps contoured at 5σ, colored by subunits (green, heavy chain; sky blue, light chain; brown, PD-1). (B) Front and back closeup views at the interface. (C) Interactions of the FG loop of PD-1 with pembrolizumab. (D) With MD-derived ED map. (E) X-ray derived ED maps contoured at 1σ superimposed onto the 5GGS crystal structures. (F) Stereo diagram of comparison of the MD-derived structure with the crystal structure. (G) A rotated view. Large displacements of K131 and Q133 of PD-1 are indicated with arrows.

MD-derived ED maps for the PD-1/pembrolizumab complex show well-defined features at the antibody–antigen interface, while peripheral regions of both proteins are less well resolved (Fig. 5). The FG loop of PD-1 forms a network of hydrogen-bonds (HBs) with Pembrolizumab: two backbone HBs with R102 of the light chain, a third backbone HB with Y53 of the heavy chain, and two side-chain HBs from K131 (a lysine finger on the FG loop) to D108 of the heavy chain and E59 of the light chain. These residues also participate in extensive intra-subunit side-chain hydrogen bonding (Fig. 5).

Notably, the interaction pattern around K131 differs between the MD-derived structure and the 5GGS crystal structure. In the MD model, R102 of the light chain forms HBs with the backbone carbonyls of K131 and Q133, whereas in the crystal structure, the interactions involve K131 and A132. Additionally, K131 forms two sidechain HBs with carboxylate groups in the MD structure, in contrast to one sidechain and one backbone carbonyl interaction in the crystal structure. These differences are accompanied by a ∼6° domain rotation of the FG loop between the two structures.

Beyond the FG loop interactions, the MD-derived PD-1/pembrolizumab complex exhibits up to 18 hydrogen bonds at the interface (see cumulative HB fractions discussed elsewhere)^28^, making pembrolizumab one of the most potent anti-PD-1 antibodies characterized to date (Fig. S1, Video 1). Notably, tyrosine residues contribute significantly to binding, with pembrolizumab providing seven Tyr residues: Y33, Y35, and Y101 from the heavy chain, and Y34, Y36, Y53, and Y57 from the light chain. These tyrosine residues form HBs with four PD-1 backbone carbonyls and three sidechains (Ser, Thr, and Lys) (Fig. S2).

Six of the 7 tyrosine sidechains of pembrolizumab display well-defined ED features, including visible central "holes" in the MD-derived ED maps, indicating limited positional fluctuation throughout the entire simulation trajectories. In addition to the large buried surface area at the interface, the MD-derived complex also exhibits a high degree of surface complementarity (Fig. S2). Such extensive and stable interactions between antigen and antibody represent a benchmark for both affinity maturation (without altering the epitope) and *de novo* antibody design (possibly for new epitopes), key goals in therapeutic antibody development.

It may be possible to reengineer this antibody using new, reliable structural information derived from MD simulations of the PD-1/pembrolizumab complex. To explore this, we conducted affinity maturation studies in parallel, starting from both the crystal structure and the MD-derived equilibrium structure of the complex.^28^ Notably, the success rate was significantly higher when using the MD-derived structure, likely reflecting its greater accuracy compared to the static crystal structure. It should be noted that the 5GGS pembrolizumab/PD-1 crystal structure is one of very few pM-affinity antibody/protein antigen complexes of known crystal structures in the PDB, including the omalizumab-IgE complex (230 pM),^80^ the dupilumab-IL4Ra complex (33-46 pM).^81^

The two PD-1/pembrolizumab complexes present in the 5GGS crystal structure show notable structural differences between the two complexes (Fig. S3).^54^ In the crystal lattice, the N-terminus of one PD-1 molecule interacts with the C-terminus of another, influencing the conformation of sidechains involved in FG loop interactions. In contrast, the MD-derived complex represents an equilibrated average structure. This distinction may help explain why structure-based antibody design efforts based solely on crystal structures often fail since crystal structures do not always represent the true solution-state conformations. MD simulations are therefore essential to distinguish between lattice-induced distortions and the configurations of the complex at thermal equilibrium in solution.

Our design process generates humanized antibodies with interfaces similar to those of naturally occurring antibodies, but quite different from typical non-antibody protein–protein interfaces. For instance, non-antibody interfaces often feature buried hydrophobic residues such as phenylalanine and other aromatics. In contrast, well-behaved therapeutic antibodies generally avoid incorporating phenylalanine at the binding site, as its high hydrophobicity can compromise solubility and promote self-aggregation or misfolding, leading to poor expression and potential toxicity. As a result, antibody–antigen interactions are often stabilized predominantly by hydrogen-bonding rather than extensive hydrophobic packing. Tyrosine is a notable exception; it is frequently found at antibody binding sites due to its ability to both form HBs and participate in limited hydrophobic interactions without substantially compromising solubility. In fact, as mentioned earlier, pembrolizumab has seven tyrosine residues at its interface with PD-1 (Fig. S2). The importance of tyrosine residues in antibodies has long been recognized.^82, 83^ This information has been implicitly included in our study when trained with the Feldhaus antibody library.^22^

Due to the distinct composition of buried residues at antibody-antigen interface, compared to non-antibody protein-protein interfaces, the approaches used for energetic analysis differ significantly. For example, the seven tyrosine residues of pembrolizumab involved in PD-1 binding primarily interact with backbone atoms rather than sidechains. When sidechain interactions do occur, they often involve small or polar residues such as Ser, Thr, Gly, and Ala. In general, antigenic epitopes tend to consist of flexible loops enriched in small or hydrophilic residues, with few buried hydrophobic residues (if any). A recurring interaction motif observed is the presence of “arginine fingers” or “lysine fingers,” where long aliphatic sidechains of antigen residues form extensive hydrophobic contacts with the antibody surface, often capped or anchored by strong terminal hydrogen-bonds. It has been long recognized that arginine and tyrosine are two most dominant (plus other two residues of serine and glycine) in all naturally occurring antibodies.^84^ The four preferred residues in naturally antibodies differ from the four residues (Tyr, Ala, Asp, and Ser) used in synthetic antibodies.^83^

Another critical feature is the clustering of multiple HBs within a localized region. As noted above, the central D122–Y68 hydrogen bond in the PD-1/PD-L1 interface contributes substantially more to binding energy than the peripheral R125-E136 salt bridge (Fig. 5), illustrating the asymmetric energetic contributions across the interface. This cluster could enhance cooperativity in contribution of binding energy and affinity.

Simple enumeration of HBs is insufficient for accurate energetic analysis. Antibody-antigen interactions involve a complex trade-off between the entropic penalty (−TΔS) associated with forming HBs at the interface and the loss of HBs with surrounding solvent molecules. Additionally, the conversion factor between buried surface area and binding free energy for antibodies differs significantly from that of non-antibody protein-protein interfaces. These factors contribute to the inherent difficulty in reliably predicting antibody-antigen interactions. As demonstrated in this study, advanced machine learning techniques can provide valuable predictions for antibody-antigen binding.

Historically, random mutagenesis of CDR loops rarely result in affinity muturation. Many resulting variants lack antibody-like biophysical properties and can exhibit undesirable traits, including aggregation, insolubility, and toxicity.^85, 86^ This high failure rate has recently led antibody engineers to adopt machine learning approaches in an effort to uncover underlying principles that govern successful antibody design.^23, 87^ One notable example is the concept of antibody "medicine-likeness," which has emerged as a valuable derived criterion for evaluating developability.^22^

### MD Simulations of six designed anti-PD-1 antibodies based on the PD-1/pembrolizumab complex

Due to the distinct interactions and residue composition at antibody-antigen interfaces compared to other protein-protein interfaces, accurate structure prediction with and without binding to an antigen remains highly challenging.^88, 89^ Notably, the six hypervariable CDR loops can adopt different conformations upon antigen binding. Beyond available tools for antibody and antibody-antigen structure prediction,^40, 90–95^ we employed MD simulations to provide information on equilibrium structures of our designed anti-PD-1 antibodies (P2N_1 to P2N_6), using the PD-1/Pembrolizumab crystal pose as a reference.^54^ Our goal was to determine whether stable complex structures could be reached as engineered from a reference complex within reasonable time scale of MD simulations.

For P2N_1 to P2N_3, the MD-derived ED maps are well defined, particularly at the subunit interface (Figs. 6–9, S4– S6). In contrast, P2N_4 to P2N_6 showed poorer convergence, with less well-resolved ED features. This does not necessarily indicate non-binding; rather, it suggests that the high-affinity binding site on PD-1 may be too distant from the initial pose for full convergence within the 400 ns MD simulation window. Indeed, we later show that starting from the PD-1/nivolumab pose, P2N_6 converges to a configuration with stronger interactions than nivolumab itself whereas the PD-1 complexes with either of P1N to P3N failed to converge when starting with the PD-1/ pembrolizumab pose.

**Figure 6.**
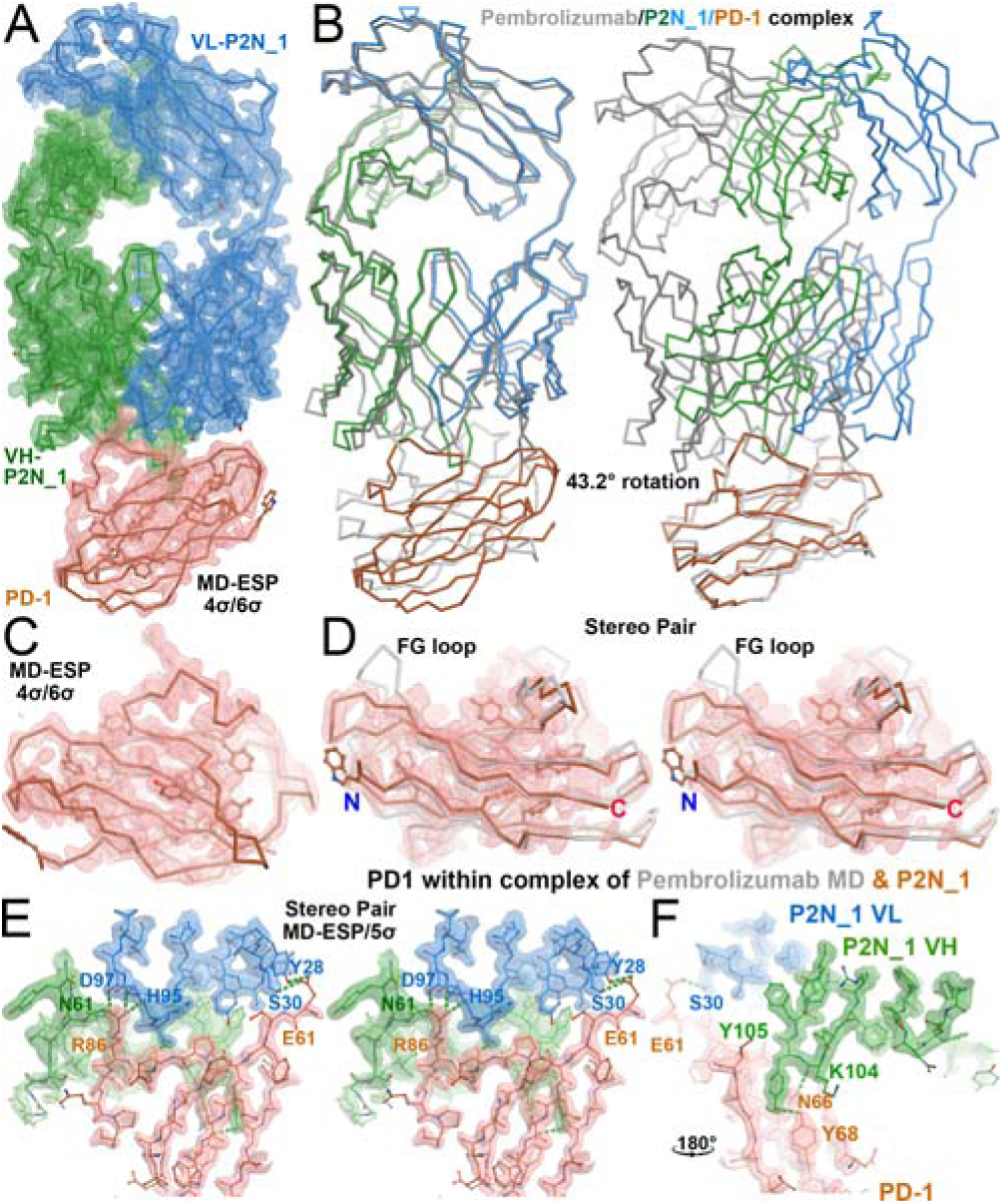
MD-derived structure for the designed P2N_1/PD-1 complex. (A) An overview with MD-derived ED map contoured at 5σ. (B) Comparison of the MD-derived equilibrium structure with the 5GGS crystal structure (silver, grey, as a starting point). Aligning either the antibody (left panel) or PD-1 (right) shows that there is a 43.2° domain rotation between them. The complex is colored by chains: light chain in sky blue, heavy chain in forest green, and PD-1and in brown. (C) An end-on view of the antibody-binding cleft of PD-1. (D) Stereo pair of PD-1 between the MD-derived (brown) and crystal structures (silver). FG loop, N and C termini differs between MD-derived and crystal structures and they are labeled. (E) Stereo pair of the inter-subunit interactions centered at an arginine finger R86 of PD-1 that forms HBs with the D97 sidechain and the H95 backbone of light chain and with the N61 sidechain of heavy chain. (F) Interaction between Y105 of heavy chain and Y68 of PD-1.

**Figure 7.**
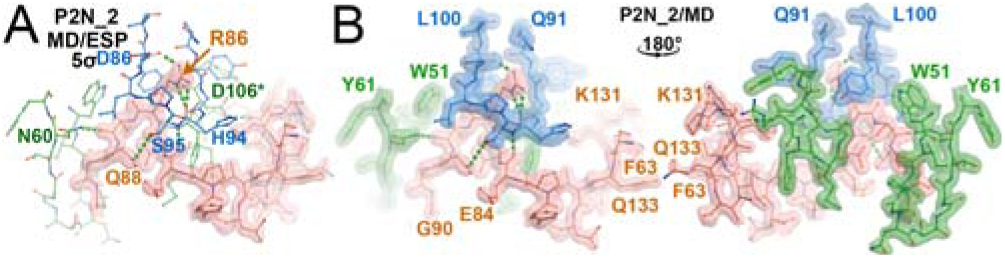
A different type of an arginine figure R86 of PD-1 within the MD-derived structure of the P2N_2/PD-1 complex in which R86 inserts into hole generated between Q91 and L100 of light chain loop of P2N_2. (A) MD-derived ED map contoured at 5σ, only for PD-1. (B) Back and front views of MD-derived map for all subunits.

**Figure 8.**
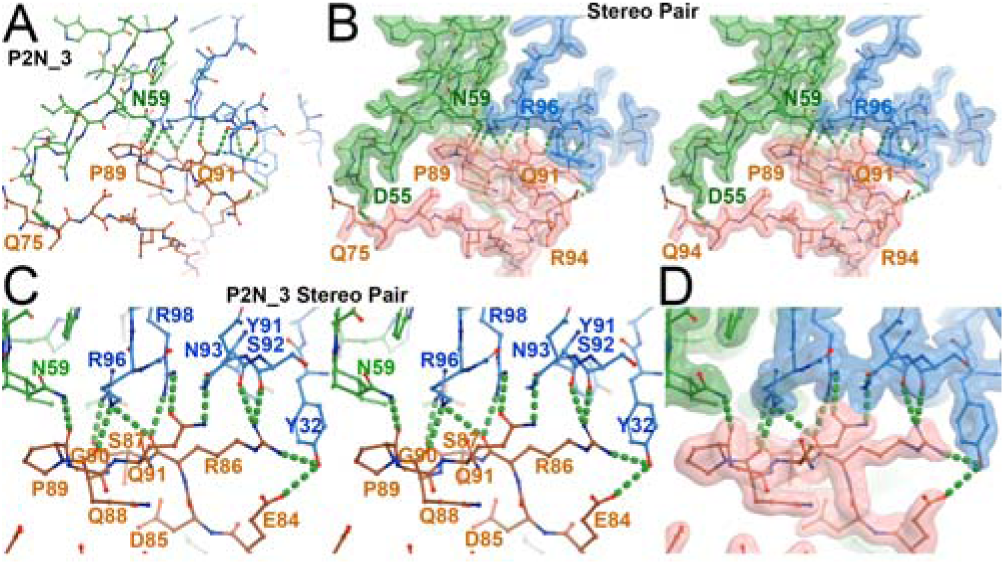
MD-derived structure for the P2N_3/PD-1 complex shows extensive HBs at the subunit interface. (A) There are 12 HBs between P2N-3 and PD-1 within the Q75 to Q91 stretch of PD-1. (B) Stereo pair with MD-derived ESP map contoured at 5σ. (C) Stereodiagram of close-up view with 11 HBs with the E84 to Q91 stretch of PD-1. (D) The corresponding MD-derived ESP map.

**Figure 9.**
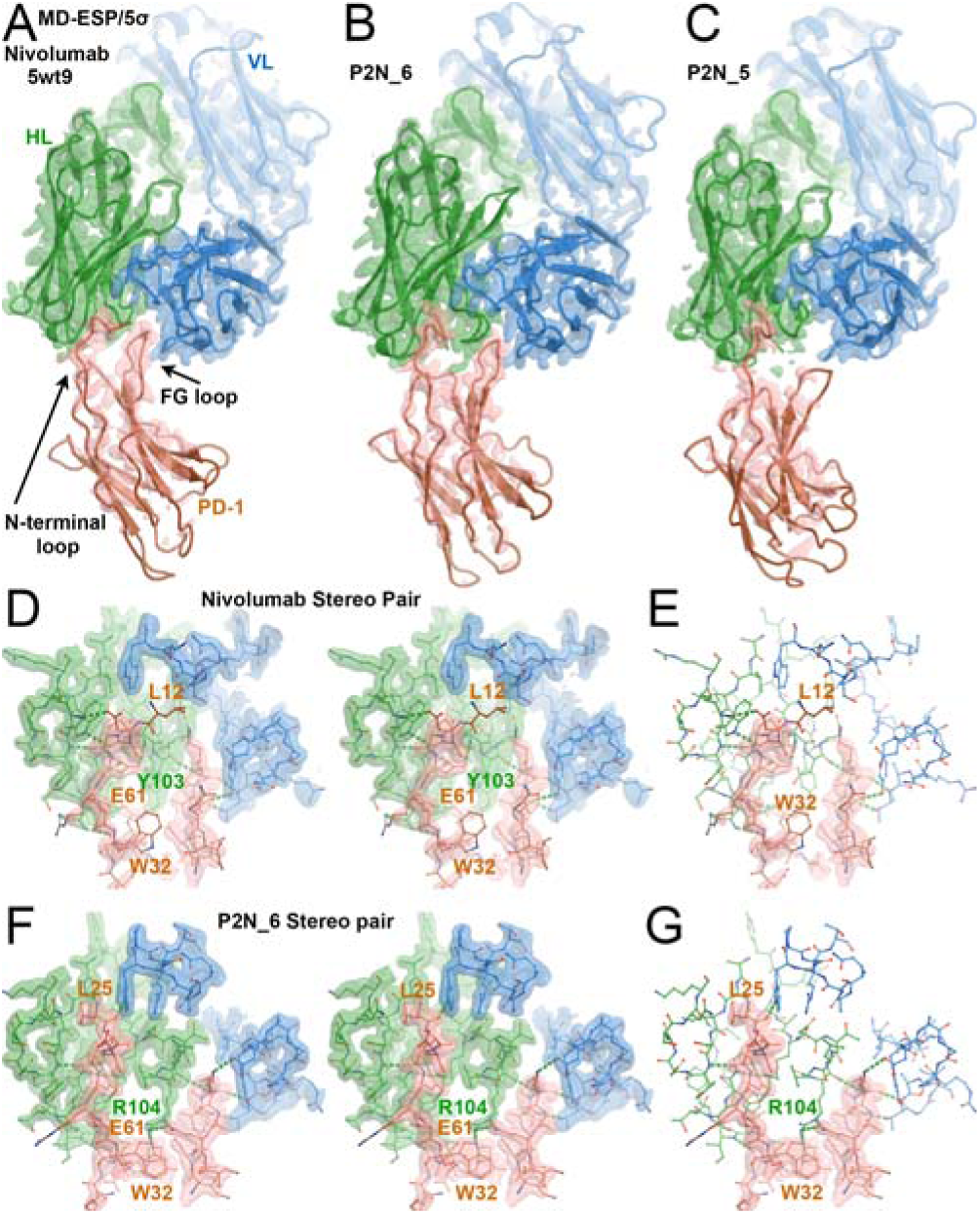
MD simulations of PD-1 complexes with nivolumab and two designed anti-PD-1 antibodies P2N_6 and P2N_5 starting with the Nivolumab pose. (A) Nivolumab with MD-derived ED map contoured at 5σ. The FG loop and N-terminal loop are labeled. (B) P2N_6. (C) P2N_6. (D) Stereodiagram of interfaces for the nivolumab/PD-1 complex superimposed onto MD-derived ED maps. (E) Omitting MD-derived map for nivolumab. (F) Stereodiagram of the P2N_6/PD-1 complex. (G) Omitting MD-derived maps for the designed antibody P2N_6.

A key indicator of stable antibody–PD-1 complex formation is the presence of well-defined features in the MD-derived ED maps for all buried interface residues, showing minimal fluctuations throughout the simulations (Figs. 5–9). Two additional structural hallmarks of stable complexes are (1) Arg/Lys "finger"-like insertions and (2) clustered hydrogen-bonds at the interface core as discussed above from the PD-1/pembrolizumab complex.

In the PD-1/pembrolizumab complex, the K131 sidechain of PD-1 inserts into a crevice formed by the heavy and light chains of the antibody (Fig. 5). Similarly, in P2N_1 and P2N_2, the R86 sidechain of PD-1 penetrates analogous structural pockets between the heavy and light chains in P2N_1 (Fig. 6), and within a pocket formed by the Q91 and L100 loops of the light chain in P2N_2 (Fig. 7). In all cases, these Arg/Lys fingers are stabilized by surrounding hydrophobic interactions along their aliphatic chains and capped by strong hydrogen-bonds of the terminal amine groups. In P2N_3, structural stability arises from a dense network of 11 inter-subunit hydrogen bonds concentrated in a short segment of PD-1 (E84–Q91), with about half of residues involving backbone carbonyl groups (Fig. 8).

For the remaining three designed anti-PD-1 antibodies, the MD-derived ED maps are relatively poorly defined, suggesting that the simulations failed to converge to stable complexes near the pembrolizumab binding pose. Among these three, the quality of ED maps follows the order: P2N_5 > P2N_6 > P2N_4. However, this does not rule out the existence of stable complexes, where alternative starting poses or different PD-1 epitopes may be required.

Another important observation is that proximity in chemical latent space does not correlate with binding orientation, binding mode, or binding affinity to PD-1 as we initially anticipated. For instance, while P2N_1 is closest to pembrolizumab in latent space, its binding orientation differs by 43.2° (Fig. 7). P2N_2, though farther in latent space, shows a smaller orientation difference of 28.4° (Figs. 7, S4), with a 24.9° difference between P2N_1 and P2N_2 (Fig. S5). These results suggest that accurate prediction of binding orientation may require AI models that incorporate antigen or epitope context. In particular, we anticipate that features such as Arg/Lys "fingers" and antigen surface fingerprints could be key elements of next-generation predictive models.^34, 96^

### MD simulations of PD-1 complexes with nivolumab and designed antibodies from the nivolumab pose

Our MD simulations of the nivolumab/PD-1 complex, initiated from the 5wt9 crystal structure, closely reproduced the experimental conformation (Fig. 9A).^55^ For this study, we used a non-glycosylated form of PD-1 at N58, which led to a minor domain rotation of 8.4° relative to the crystal structure. This shift slightly reoriented both the FG loop and the glycosylation loop. Such domain flexibility is allowed in MD simulations but is likely constrained by crystal lattice contacts in the experimental structure.

The interaction pattern between nivolumab and PD-1 is largely preserved in both the MD-derived equilibrium structure and the crystal structure, particularly involving the FG and N-terminal loops (Fig. 9A). A key exception is near the glycosylation site: in the crystal structure, PD-1 residue E61 forms three HBs with Y61 and T23 sidechains and the backbone amide of Y23 in the nivolumab heavy chain. In the MD simulations, E61 instead points toward the glycosylation pocket. A similar recognition pattern is seen in the designed antibodies P2N_6 and P2N_5 (Fig. 9), with P2N_6 sharing the same latent chemical space position as nivolumab.

Based on the quality of MD-derived ED maps at the antibody/PD-1 interface, the complex stability follows the following order: P2N_6 > nivolumab > P2N_5. In the P2N_6 complex, PD-1 residues L25 and W32 are well ordered and clearly defined in the equilibrium structure. Heavy chain R104 of P2N_6 lies within hydrogen-bonding distance of W32, which itself is H-bonded to Q133 of PD-1. This R104 is absent in both nivolumab and P2N_5. PD-1 L25 forms a backbone hydrogen bond with heavy chain W94 of P2N_6, and its sidechain is buried in a hydrophobic pocket. W94 is stacked against R96 of the same chain, which forms three HBs: one with the N93 backbone and two with D100 of the light chain. While W94 and R96 are conserved in nivolumab, their conformations differ significantly.

### Hydrogen bonding probability in MD trajectories

The R86 “finger” of PD-1 is recognized differently by P2N_1 and P2N_2 (Fig. 10). In P2N_2, R86 inserts into a compact pocket formed by residues Y92 to D98 within the Q91–L100 loop region (Figs. 7, 10A). The MD-derived ED map reveals that the equilibrium structure exhibits a dominant conformation, where R86Nε and R86Nη2 form hydrogen-bonds with the H94 carbonyl, while R86Nη1 forms a hydrogen-bond with the D98 carbonyl (Fig. 10).

**Figure 10.**
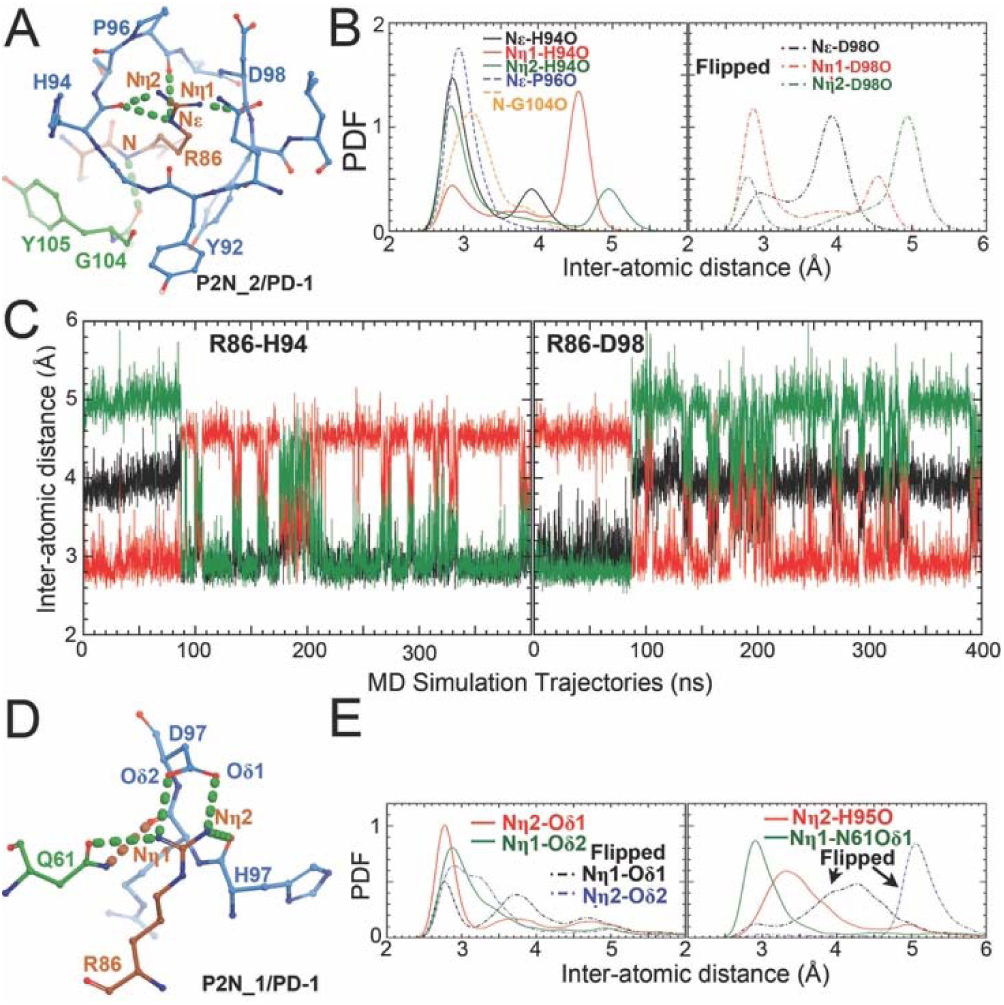
Hydrogen bonding interactions of R86 arginine fingers of PD-1 in MD simulation trajectories. (A) R86 of PD-1 in the equilibrium fitted structure within the P2N_2/PD-1 complex. (B) Probability distribution function (PDF) distribution of inter-atomic distances for HBs (left panel) and flipped R86 sidechain (right panel). (C) Inter-atomic distance trajectories as a function of time. (D) R86 of PD-1 in the equilibrium fitted structure within the P2N_1/PD-1 complex. (E) PDF distributions of inter-atomic distances.

The MD trajectory analysis reveals a minor conformation where the R86 sidechain is flipped. The minor conformation dominated the first 100 ns of simulation, after which it was replaced by the more stable, dominant state for the remaining 300 ns. During this period, only one sidechain flip was observed among 10 possible attempts. Similar analyses have previously uncovered hidden states in other systems (e.g., the HNH domain of CRISPR enzyme).^79, 97^

Residue R86 of PD-1 forms four hydrogen bonds with P2N_1. However, the paired probability distribution function (PDF) for this complex is significantly broader than that observed in the P2N_2/PD-1 complex. While the P2N_2/PD-1 system exhibits a bimodal PDF distribution, the P2N_1/PD-1 complex displays a complex, asymmetric tailing without distinct peaks. These interatomic PDFs reflect the underlying free energy landscape sampled during MD simulations.

In the P2N_2/PD-1 complex, the R86 sidechain exhibits minimal conformational fluctuations, indicating a stable configuration. Larger fluctuations are not favorable and primarily occur when the sidechain attempts to transition from the dominant conformational state to a minor one. In contrast, the broader PDF observed in the P2N_1/PD-1 complex suggests that the HB network involving R86 is less constrained, and exhibits greater flexibility and conformational sampling.

The PDF of HB distances shows that HBs formed between PD-1 and P2N_3 have more pronounced peaks in PDF than those in complexes with P2N_2 or P2N_1 (Figs. 10, 11). These strong HBs effectively constrain the conformational flexibility of key sidechains, preventing flipping of arginine and glutamine residues throughout the MD simulations—except for the E84 sidechain of PD-1, which displays rapid flipping as shown by weak electrostatic potential (ESP) features (Fig. 11).

**Figure 11.**
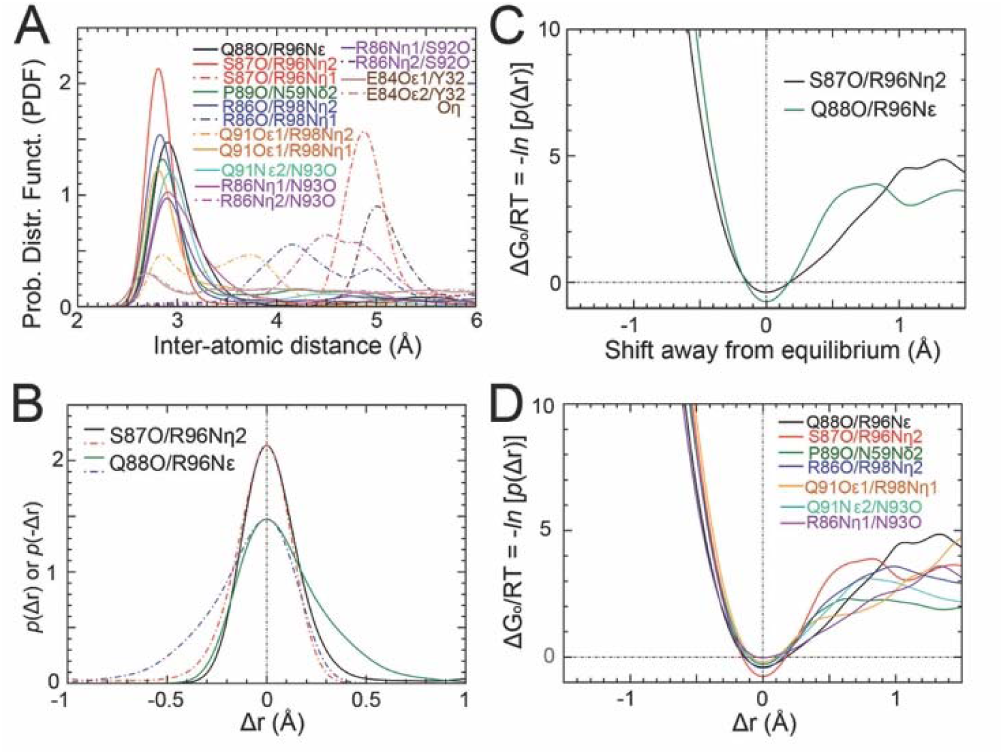
Local free energy landscape of hydrogen bonding interactions. (A) *P*(*r*) PDF for HBs between PD-1 and P2N_3. Flipped geometry is shown in dashed curves. (B) Two examples of P(r) being converted their *P*(Δr) where Δr is a deviation to the most equilibrium distance. (C) Converting PDF to free energy through the ΔG_o_/RT = -*ln* [*P*(Δr)] relationship. (D) For all major HBs between PD-1 and P2N_3.

The PDF of HB distances typically follow the Boltzmann distribution, -*ln* [*P*(Δr)] = ΔG_0_/RT, with a higher peaks corresponding to more stable free energy configurations. For instance, the HB between S87O (PD-1) and R98 (P2N_3) defines a configuration approximately 4Qtimes RT deeper than the Q88 (PD-1)/R98 (P2N_3) interaction (Fig. 11C). The total free energy contribution due to HBs between PD-1 and P2N_3 is estimated to be ∼30QRT units, providing exceptional stability of the complex. This stability may even surpass that of the parental pembrolizumab/PD-1 complex from which the antibodies were derived.

#### Binding kinetics of PD-L1 and designed antibodies to PD-1

With the exception of P2N_1, all five designed anti-PD-1 antibodies, along with pembrolizumab, nivolumab (in both Fab fragment and full IgG formats), PD-1, and PD-L1 expressed in Expi293F cells were successfully and readily purified to homogeneity (Fig. S10). Binding experiments between PD-1 and either PD-L1 or PD-L2 were conducted at a concentration of 1QμM. The association kinetics revealed a predominant fast-binding phase, comprising 75.5% and 90.0% of the population for PD-L1 and PD-L2, respectively (Fig. 12). A slower association phase accounted for 34.7% of the population for PD-L1 and 50.1% for PD-L2. The slowest dissociation component, likely reflecting baseline drift, was observed with a minor population ranging from 4.3% to 12.0%. The concentration we selected in this study was try to balance simultaneously for both high-affinity and low-affinity binding complexes, and it not optimal for very high and very low affinity complexes.

**Figure 12.**
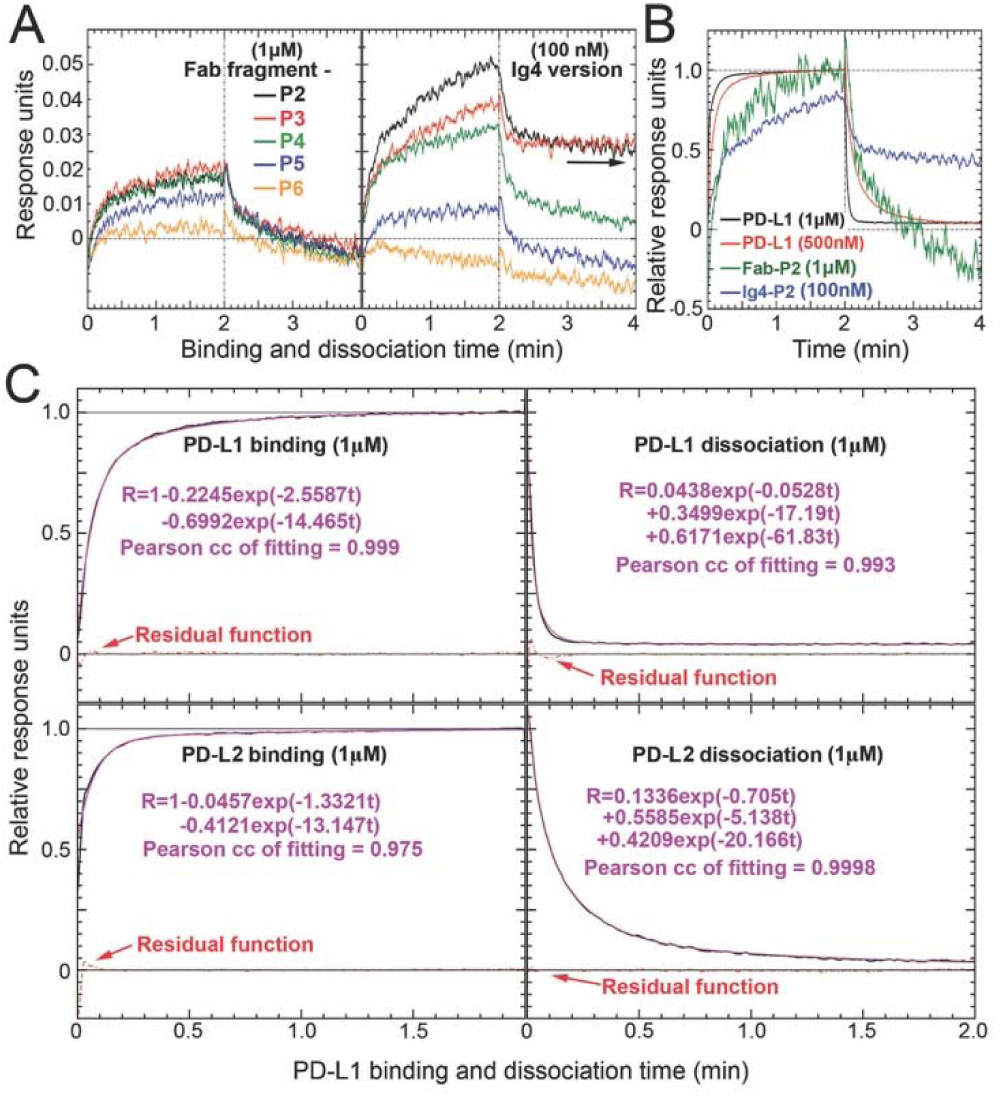
Association/dissociation phases of kinetic studies of designed antibodies, PD-L1 and PD-L2 to PD-1. (A) Binding of designed P2, P3, P4, P5, and P6 fragments at 1 μM (left panels) or IgG versions at 100 nM (right). Arrow indicates large fraction of the formed complex remains for P2 and P3 Ig versions at the 2 minute end of dissociation phase. (B) Comparison of binding of P2 fragment at 1 μM (green curve), P2 IgG at 100 nM (blue) with binding of PD-1 (black) and PD-L2 (red) at 1 μM after normalization by making the asymptotic values being the unity. (C) Fitting multiple-exponential functions of PD-L1 (upper panels) and PD-L2 (lower panels) binding to PD-1 at both 1 μM for both association (left panels) and dissociation panel (right). Both Goodness of fit using Pearson correlation coefficients and residual functions (red) are included.

The binding affinity, calculated as *K*_D_ = *k*_off_/*k*_on_ = [*k*_off_/(*k*_fast_+*k*_off_)][Antibody]_0_, was determined using fitted parameters from the fast association and slow dissociation phases. The resulting *K*_D_ values ranged from 0.54 to 0.81 μM for PD-L1 and from 0.14 to 0.30 μM for PD-L2. These values indicate stronger binding affinities in our study than values previously reported, ranging from 2 to 8 μM for PD-L1, and 0.5 to 2.1 μM for PD-L2.^98–101^ Some of these discrepancies can be explained by the fact that many of earlier studies assumed that there was a single-binding phase in complex formation.

With the exception of the P2N_6 variant, four of the five purified antibody variants exhibited classical association profiles, although their response signals were significantly lower than those observed for the PD-1/PD-L1 complex (Fig. 12) or for pembrolizumab or nivolumab (Fig. 13). Nevertheless, the IgG versions of these variants produced slightly higher response signals compared to their Fab fragments (Fig. 12A). In the IgG format, more than 50% of the complexes formed by P2N_2 and P2N_3 remained bound at the 2-minute endpoint of the dissociation phase, indicating a half-life time exceeding 2 minutes. By contrast, the PD-1/PD-L1 complex has a dissociation half-life of approximately 3 to 15 seconds.

**Figure 13.**
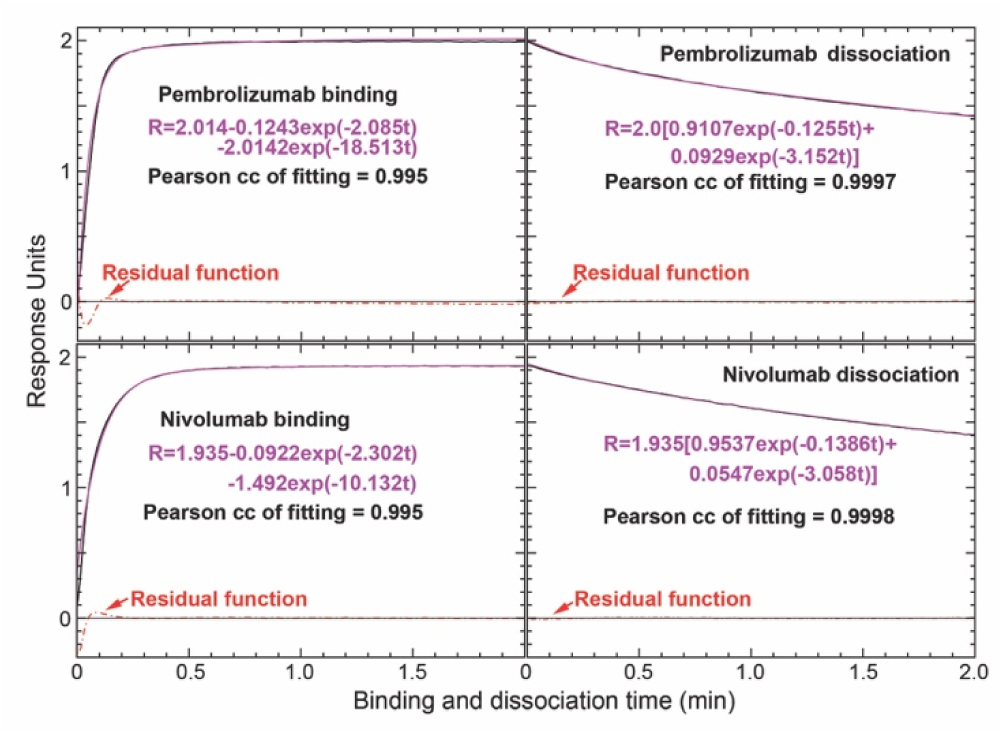
Binding assays of pembrolizumab (upper panels) or nivolumab (lower panels) Fab fragment at 1 μM to immobilized PD-1 with fitted kinetic parameters for both association (left panels) and dissociation (right panels) panels. Pearson correlation coefficients (goodness of fit) and residual functions (red dashed curves) are shown.

In the Fab versions of P2N_2 (P2), P2N_3 (P3), P2N_4 (P4), and P2N_5 (P5), the binding curves approached asymptotic phases by the 2-minute mark, whereas this behavior was not observed in the IgG versions (Fig. 12). It is unexpected that all designed antibodies have very low signals in the experiment, the reason for which remains unknown. One possibility is that the specific tag on PD-1 we used may somehow interfere with the binding assay. Due to the low signal intensity, kinetic rate constants were not determined for individual phases of these variants. It is worth noting that some single-point mutations in human PD-1 have previously been shown to significantly alter response profiles (also with much reduced signal intensities) in similar PD-L1 binding assays using BLI.^101^ However, the effects observed here in our designed antibodies were substantially greater than those reported previously for such PD-1 point mutations.

In parallel, binding assays were performed for pembrolizumab and nivolumab in both Fab fragment form (at 1QμM; Fig. 13) and IgG format (at 100QnM; data not shown). For the Fab fragments, the determined equilibrium dissociation constant (K_D_) was 3.2QnM for pembrolizumab and 6.7QnM for nivolumab. These values suggest a much smaller difference in binding affinity between the two antibodies than previously reported, where K_D_ values of 29QpM for pembrolizumab and 4.0QnM for nivolumab have been documented.^55, 56, 102^ However, it should be noted that the kinetic parameters derived from the fast association phase in our current assays are only approximately determined due to the instrument dead time at high concentration conditions (Fig. 13).

For the IgG versions of pembrolizumab and nivolumab tested at 100 nM, the binding kinetics did not reach asymptotic values within the 2-minute observation window, and as a result, no kinetic fitting was performed. In a separate study involving these antibodies and a different set of designed variants at a concentration of 200QnM, the binding curves were more complete and suitable for accurate kinetic analysis, as reported elsewhere.^28^

Although binding affinities for the designed anti-PD-1 antibodies could only be estimated under the current experimental conditions, the overall binding trends align with predictions from MD simulations. Specifically, variants P2N_2 and P2N_3 are predicted to exhibit stronger binding interactions compared to P2N_5 and P2N_6.

It remains unclear why the BLI response signals for binding and dissociation of our designed antibodies to immobilized PD-1 are so weak when compared with those of the parental pembrolizumab and its computational affinity maturated antibodies.^28^ It remains to be addressed whether particular immobilization of PD-1 could have somehow impaired the binding kinetics because our designed antibodies recognize very different epitopes from the parental pembrolizumab. It should be noted that our MD simulations were carried out at physiologically relevant temperature of 37°C while the binding assays were done at room temperature of 25°C. We have not yet examined the issues of temperature-dependent binding assays of either the parental or designed antibodies to PD-1, nor as to whether designed antibodies have been post-translationally modified in an unexpected manner.

Interestingly, the reported asymptotic response units of fragment-based computation design of single-body antibodies against human serum albumin (HSA) is also about 0.015 units using the same BLI methodology,^47^ values similar to ours. In fact, up to date, almost all *de nova* designed antibodies started with relatively low affinity, including those derived using Rosetta Fold-based diffusion (RFDiffusion) method, and much higher affinity antibodies were obtained only after experimental affinity maturation using the OrthoRep method.^103^ OrthoRep is a laboratory-evolutionary system with enhanced mutational rate about 1 millions times faster than the naturally-occurring rate.^29, 30^ Starting with brand new antibodies discovered in this study, it might be possible to produce high affinity anti-PD-1 antibodies when combined with either computational or experimental affinity maturation methods. ^28–30^

It is noted that large variations in response units over 20 fold were recently observed for a group of redesigned anti-PD-1 mutants using a computation saturation mutational approach, starting with pembrolizumab even though the determined apparent K_D_ value varied only about 40% from 62 to 85 pM.^28^ Therefore the asymptotic response units alone does not bear any correlation with binding affinity. The dissociation constant for pembrolizumab from its PD-1 complex determined in this study was 3.2 nM (without additional concentration optimization), which differs significantly from the recently determined value of 0.042 nM.^28^ This difference highlights variations in experimental conditions such as the extent of both immobilization and concentration optimization. The concentrations of antibodies used in this study were much higher than the previous study.^28^ The effects of immobilization used in binding affinity studies are well documented, i.e., the association constant for the IL4Rα ectodomain-rademikibart complex was determined at 20.7 pM when IL4Rα was immobilized, but it was 448 pM when rademikibart was immobilized.^104^ The corresponding values were 45.8 and 516 pM for the IL4Rα/dupilumab complex.^104^

Because of these experimental limitations, we were unable at the moment to fully validate our MD simulation results that P2N_3 and P2N_6 should have better binding profiles than nivolumab to PD-1. Lastly, the antibody sequence library used for training in this study has no specific information on binding affinity (typically from μM to pM) to antigens. It remains to be seen whether traits of high-affinity antibodies differ from low-affinity ones. If so, the affinity-based selected antibodies can be used for further AI training.

## CONCLUDING REMARKS

Antibody engineering remains a significant challenge, particularly when employing randomized mutagenesis within the six complementarity-determining regions. The majority of resulting variants tend to exhibit poor biophysical properties, including aggregation, low solubility, or high cytotoxicity. Among these, only a small fraction demonstrates specific antigen binding, and even then, off-target interactions are frequently observed. To address these limitations, computational approaches, including structure-based modeling and machine learning, are being developed to better predict and guide antibody–antigen interactions.

In this study, we introduced a machine learning–driven framework that incorporates "antibody medicine-likeness" as a core design principle, with the goal of generating antibody variants that are both biophysically stable and therapeutically relevant. Using the CKEA, we generated new antibody candidates by interpolating between two FDA-approved anti–PD-1 antibodies, nivolumab and pembrolizumab, within a chemically informed latent space containing hidden information of PD-1 epitopes. This generative model was specifically tailored to preserve desirable pharmacological features while sampling variants with potential for improved binding affinity. With the exception of one designed antibody P2N_1, which has a favorable complex formation according to MD simulations but failed to be purified, all the remaining 5 antibodies have desirable medicine-likeness properties.

To assess the structural and functional viability of these designs, we employed MD simulations to model equilibrium binding conformations with the PD-1 antigen. Experimental validation included expression, purification, and binding assays for the designed variants. Notably, three of the six designed antibodies demonstrated specific binding to PD-1 with reasonable saturation curves (although signals were relatively weak for thermodynamics and kinetics analysis), approaching their asymptotic values within 2 min, which are on comparable time scale for formation of the PD-1/PD-L1 interaction. Based on the quality of MD-derived ED maps at the antibody/PD-1 interface, the complex stability follows the following order: P2N_6 > nivolumab > P2N_5.

Together, our results highlight the utility of combining AI-guided design with physics-based simulation and experimental validation in a unified pipeline for antibody discovery. This integrative approach is broadly applicable across a wide range of immunotherapeutic targets and provides a generalizable and scalable framework for the rational design of therapeutically viable antibodies.

## ASSOCIATED CONTENT

### Supporting Information

The Supporting Information is available free of charge at https:/pubs.ac.org/dox/acs.biochemistry.xxxxxxx Supporting Tables S1-S3, Supporting Figures S1-S10, Supporting Video 1

Computer folder archive containing the equilibrium atomic coordinates for 6 designed P2N_n.pdb where n =1, 2, 3, 4, 5, and 6, as well as for the PD1/PDL1 complex and PD-1/nivolumab complex and their MD-derived ED/ESP maps (ZIP). The MD-ED map for the PD-1/pembrolizumab complex coordinates were available in Supporting Information of reference 28 (Shi et al., 2026).^28^

## DATA AVAILABILITY

Pretrained KEA can be accessed through an API interface at kaemd.batistalab.com. For inquiries and instructions, please reach out to Dr. Victor S. Batista.

## AUTHOR INFORMATION

Corresponding Authors: To whom correspondence should be addressed: S.T., J.W., and V.S.B.

## Author Contributions

V.S.B., S.T., and J.W. conceived the idea and designed studies, Y. Shi., H.L. carried out studies, Y. Shi, and J.W. carried out analysis and wrote the draft manuscript with input from all authors. Y.L and S.T. for experimental validation. J.W., C. G. B., S.T., V.S.B. oversaw the entire study, edited and approved the final manuscript.

## ACKNOWLEDGMENT

Authors thank Ms. Yeil Kim for reading and commenting on the manuscript and for helping some experiments. This study was funded by the National Institutes of Health under Grants No. R01 GM136815 (V.S.B) and R00HD104924 (S.T.), Carol and Gene Ludwig Pilot Project Award (S. T.), and carried out with computation times provided by the National Research Scientific Computing Center (NERSC) under Grant No. M3807.

## ABBREVIATIONS

PD-1: Programmed cell Death protein-1 (CD279)
PD-L1: programmed cell death protein ligand-1 (CD274)
PD-L2: programmed cell death protein ligand 2 (CD273)
P2N_n: CKEA-extrapolated anti-PD-1 antagonistic antibodies number (n)
CKEA: conditional kernel-elastic autoencoder
MD: molecular dynamics
ED: electron density
HB: hydrogen bond.

## REFERENCES

(1) Lazar-Molnar, E.; Chen, B.; Sweeney, K. A.; Wang, E. J.; Liu, W.; Lin, J.; Porcelli, S. A.; Almo, S. C.; Nathenson, S. G.; Jacobs, W. R., Jr. Programmed death-1 (PD-1)-deficient mice are extraordinarily sensitive to tuberculosis. Proc Natl Acad Sci U S A 2010, 107 (30), 13402–13407. DOI: 10.1073/pnas.1007394107.

(2) Frebel, H.; Nindl, V.; Schuepbach, R. A.; Braunschweiler, T.; Richter, K.; Vogel, J.; Wagner, C. A.; Loffing-Cueni, D.; Kurrer, M.; Ludewig, B.;, et al. Programmed death 1 protects from fatal circulatory failure during systemic virus infection of mice. J Exp Med 2012, 209 (13), 2485–2499. DOI: 10.1084/jem.20121015.

(3) Ohaegbulam, K. C.; Assal, A.; Lazar-Molnar, E.; Yao, Y.; Zang, X. Human cancer immunotherapy with antibodies to the PD-1 and PD-L1 pathway. Trends Mol Med 2015, 21 (1), 24–33. DOI: 10.1016/j.molmed.2014.10.009.

(4) Philips, G. K.; Atkins, M. Therapeutic uses of anti-PD-1 and anti-PD-L1 antibodies. Int Immunol 2015, 27 (1), 39–46. DOI: 10.1093/intimm/dxu095.

(5) Balar, A. V.; Weber, J. S. PD-1 and PD-L1 antibodies in cancer: current status and future directions. Cancer Immunol Immunother 2017, 66 (5), 551–564. DOI: 10.1007/s00262-017-1954-6.

(6) Han, Y.; Liu, D.; Li, L. PD-1/PD-L1 pathway: current researches in cancer. Am J Cancer Res 2020, 10 (3), 727–742.

(7) Mueller, S. N.; Vanguri, V. K.; Ha, S. J.; West, E. E.; Keir, M. E.; Glickman, J. N.; Sharpe, A. H.; Ahmed, R. PD-L1 has distinct functions in hematopoietic and nonhematopoietic cells in regulating T cell responses during chronic infection in mice. J Clin Invest 2010, 120 (7), 2508–2515. DOI: 10.1172/JCI40040.

(8) Alsaab, H. O.; Sau, S.; Alzhrani, R.; Tatiparti, K.; Bhise, K.; Kashaw, S. K.; Iyer, A. K. PD-1 and PD-L1 Checkpoint signaling inhibition for cancer immunotherapy: Mechanism, Combinations, and Clinical Outcome. Front Pharmacol 2017, 8, 561. DOI: 10.3389/fphar.2017.00561.

(9) Weigmann, K. Releasing the brakes to fight cancer: The recent discovery of checkpoints has boosted the field of cancer immunotherapy. EMBO Rep 2016, 17 (9), 1257–1260. DOI: 10.15252/embr.201643038.

(10) Wong, R. M.; Scotland, R. R.; Lau, R. L.; Wang, C.; Korman, A. J.; Kast, W. M.; Weber, J. S. Programmed death-1 blockade enhances expansion and functional capacity of human melanoma antigen-specific CTLs. Int Immunol 2007, 19 (10), 1223–1234. DOI: 10.1093/intimm/dxm091.

(11) Wang, C.; Thudium, K. B.; Han, M.; Wang, X. T.; Huang, H.; Feingersh, D.; Garcia, C.; Wu, Y.; Kuhne, M.; Srinivasan, M.;, et al. In vitro characterization of the anti-PD-1 antibody nivolumab, BMS-936558, and in vivo toxicology in non-human primates. Cancer Immunol Res 2014, 2 (9), 846–856. DOI: 10.1158/2326-6066.CIR-14-0040.

(12) Rajan, A.; Gulley, J. L. Nivolumab (anti-PD-1, BMS-936558, ONO-4538) in patients with advanced non-small cell lung cancer. Transl Lung Cancer Res 2014, 3 (6), 403–405. DOI: 10.3978/j.issn.2218-6751.2014.09.02.

(13) Hamid, O.; Robert, C.; Daud, A.; Hodi, F. S.; Hwu, W. J.; Kefford, R.; Wolchok, J. D.; Hersey, P.; Joseph, R. W.; Weber, J. S.;, et al. Safety and tumor responses with lambrolizumab (anti-PD-1) in melanoma. N Engl J Med 2013, 369 (2), 134–144. DOI: 10.1056/NEJMoa1305133.

(14) Robert, C.; Ribas, A.; Wolchok, J. D.; Hodi, F. S.; Hamid, O.; Kefford, R.; Weber, J. S.; Joshua, A. M.; Hwu, W. J.; Gangadhar, T. C.;, et al. Anti-programmed-death-receptor-1 treatment with pembrolizumab in ipilimumab-refractory advanced melanoma: a randomised dose-comparison cohort of a phase 1 trial. Lancet 2014, 384 (9948), 1109–1117. DOI: 10.1016/S0140-6736(14)60958-2.

(15) Booth, B. J.; Ramakrishnan, B.; Narayan, K.; Wollacott, A. M.; Babcock, G. J.; Shriver, Z.; Viswanathan, K. Extending human IgG half-life using structure-guided design. MAbs 2018, 10 (7), 1098–1110. DOI: 10.1080/19420862.2018.1490119.

(16) Ko, S.; Jo, M.; Jung, S. T. Recent achievements and challenges in prolonging the serum half-lives of therapeutic IgG antibodies through Fc Engineering. BioDrugs 2021, 35 (2), 147–157. DOI: 10.1007/s40259-021-00471-0.

(17) Li, H.; Shee, Y.; Allen, B.; Maschietto, F.; Morgunov, A.; Batista, V. S. Kernel-elastic autoencoder for molecular design. PNAS Nexus 2024, 3, pgae168. DOI: https://academic.oup.com/pnasnexus/article/3/4/pgae168/7658379.

(18) Cuomo, A. E.; Ibarraran, S.; Sreekumar, S.; Li, H.; Eun, J.; Menzel, J. P.; Zhang, P.; Buono, F.; Song, J. J.; Crabtree, R. H.;, et al. Feed-forward neural network for predicting enantioselectivity of the asymmetric negishi reaction. ACS Cent Sci 2023, 9 (9), 1768–1774. DOI: 10.1021/acscentsci.3c00512.

(19) Chames, P.; Van Regenmortel, M.; Weiss, E.; Baty, D. Therapeutic antibodies: successes, limitations and hopes for the future. Br J Pharmacol 2009, 157 (2), 220–233. DOI: 10.1111/j.1476-5381.2009.00190.x.

(20) Lu, R. M.; Hwang, Y. C.; Liu, I. J.; Lee, C. C.; Tsai, H. Z.; Li, H. J.; Wu, H. C. Development of therapeutic antibodies for the treatment of diseases. J Biomed Sci 2020, 27 (1), 1. DOI: 10.1186/s12929-019-0592-z.

(21) Carter, P. J.; Rajpal, A. Designing antibodies as therapeutics. Cell 2022, 185 (15), 2789–2805. DOI: 10.1016/j.cell.2022.05.029.

(22) Ahmed, L.; Gupta, P.; Martin, K. P.; Scheer, J. M.; Nixon, A. E.; Kumar, S. Intrinsic physicochemical profile of marketed antibody-based biotherapeutics. Proc Natl Acad Sci U S A 2021, 118 (37). DOI: 10.1073/pnas.2020577118.

(23) Makowski, E. K.; Kinnunen, P. C.; Huang, J.; Wu, L. N.; Smith, M. D.; Wang, T. X.; Desai, A. A.; Streu, C. N.; Zhang, Y. L.; Zupancic, J. M.;, et al. Co-optimization of therapeutic antibody affinity and specificity using machine learning models that generalize to novel mutational space. Nature Communications 2022, 13 (1). DOI: ARTN 3788 https://10.1038/s41467-022-31457-3.

(24) de Wildt, R. M.; Mundy, C. R.; Gorick, B. D.; Tomlinson, I. M. Antibody arrays for high-throughput screening of antibody-antigen interactions. Nat Biotechnol 2000, 18 (9), 989–994. DOI: 10.1038/79494.

(25) Holt, L. J.; Bussow, K.; Walter, G.; Tomlinson, I. M. By-passing selection: direct screening for antibody-antigen interactions using protein arrays. Nucleic Acids Res 2000, 28 (15), E72. DOI: 10.1093/nar/28.15.e72.

(26) Makowski, E. K.; Chen, H. W.; Lambert, M.; Bennett, E. M.; Eschmann, N. S.; Zhang, Y. L.; Zupancic, J. M.; Desai, A. A.; Smith, M. D.; Lou, W. J.;, et al. Reduction of therapeutic antibody self-association using yeast-display selections and machine learning. Mabs 2022, 14 (1). DOI: Artn 2146629 https://10.1080/19420862.2022.2146629.

(27) Starr, C. G.; Makowski, E. K.; Wu, L. N.; Berg, B.; Kingsbury, J. S.; Gokarn, Y. R.; Tessier, P. M. Ultradilute measurements of self-association for the identification of antibodies with favorable high-concentration solution properties. Mol Pharmaceut 2021, 18 (7), 2744–2753. DOI: 10.1021/acs.molpharmaceut.1c00280.

(28) Shi, Y.; Kim, Y.; Liu, P.; Wang, J.; Tang, S.; Batista, V. S. Computational evolution of anti-PD-1 antibodies induces structural refolding for high-affinity interactions. Biochemistry 2026, 65, 517–520. DOI: 10.1021/acs.biochem.5c00574.

(29) Rix, G.; Watkins-Dulaney, E. J.; Almhjell, P. J.; Boville, C. E.; Arnold, F. H.; Liu, C. C. Scalable continuous evolution for the generation of diverse enzyme variants encompassing promiscuous activities. Nat Commun 2020, 11 (1), 5644. DOI: 10.1038/s41467-020-19539-6.

(30) Rix, G.; Williams, R. L.; Hu, V. J.; Spinner, A.; Pisera, A. O.; Marks, D. S.; Liu, C. C. Continuous evolution of user-defined genes at 1 million times the genomic mutation rate. Science 2024, 386 (6722), eadm9073. DOI: 10.1126/science.adm9073.

(31) Lippert, A. H.; Paluch, C.; Gaglioni, M.; Vuong, M. T.; McColl, J.; Jenkins, E.; Fellermeyer, M.; Clarke, J.; Sharma, S.; Moreira da Silva, S.;, et al. Antibody agonists trigger immune receptor signaling through local exclusion of receptor-type protein tyrosine phosphatases. Immunity 2024, 57 (2), 256–270 e210. DOI: 10.1016/j.immuni.2024.01.007.

(32) Liu, L.; Takashima, S.; Tokumaru, Y.; Ikuta, N.; Nakajima, Y.; Ohta, A. Identification of anti-mouse PD-1 agonist antibodies that inhibit T cell activation. Front Immunol 2025, 16, 1631929. DOI: 10.3389/fimmu.2025.1631929.

(33) Suzuki, K.; Tajima, M.; Tokumaru, Y.; Oshiro, Y.; Nagata, S.; Kamada, H.; Kihara, M.; Nakano, K.; Honjo, T.; Ohta, A. Anti-PD-1 antibodies recognizing the membrane-proximal region are PD-1 agonists that can down-regulate inflammatory diseases. Sci Immunol 2023, 8 (79), eadd4947. DOI: 10.1126/sciimmunol.add4947.

(34) Gainza, P.; Sverrisson, F.; Monti, F.; Rodola, E.; Boscaini, D.; Bronstein, M. M.; Correia, B. E. Deciphering interaction fingerprints from protein molecular surfaces using geometric deep learning. Nat Methods 2020, 17 (2), 184–192. DOI: 10.1038/s41592-019-0666-6.

(35) Marchand, A.; Buckley, S.; Schneuing, A.; Pacesa, M.; Elia, M.; Gainza, P.; Elizarova, E.; Neeser, R. M.; Lee, P. W.; Reymond, L.;, et al. Targeting protein-ligand neosurfaces with a generalizable deep learning tool. Nature 2025, 639 (8054), 522–531. DOI: 10.1038/s41586-024-08435-4.

(36) Mason, D. M.; Friedensohn, S.; Weber, C. R.; Jordi, C.; Wagner, B.; Meng, S. M.; Ehling, R. A.; Bonati, L.; Dahinden, J.; Gainza, P.;, et al. Optimization of therapeutic antibodies by predicting antigen specificity from antibody sequence via deep learning. Nat Biomed Eng 2021, 5 (6), 600–612. DOI: 10.1038/s41551-021-00699-9.

(37) Yoshida, K.; Kuroda, D.; Kiyoshi, M.; Nakakido, M.; Nagatoishi, S.; Soga, S.; Shirai, H.; Tsumoto, K. Exploring designability of electrostatic complementarity at an antigen-antibody interface directed by mutagenesis, biophysical analysis, and molecular dynamics simulations. Sci Rep 2019, 9 (1), 4482. DOI: 10.1038/s41598-019-40461-5.

(38) Di Rienzo, L.; Milanetti, E.; Lepore, R.; Olimpieri, P. P.; Tramontano, A. Superposition-free comparison and clustering of antibody binding sites: implications for the prediction of the nature of their antigen. Sci Rep 2017, 7, 45053. DOI: 10.1038/srep45053.

(39) Di Rienzo, L.; Milanetti, E.; Ruocco, G.; Lepore, R. Quantitative description of surface complementarity of antibody-antigen interfaces. Front Mol Biosci 2021, 8, 749784. DOI: 10.3389/fmolb.2021.749784.

(40) Ruffolo, J. A.; Chu, L. S.; Mahajan, S. P.; Gray, J. J. Fast, accurate antibody structure prediction from deep learning on massive set of natural antibodies. Nat Commun 2023, 14 (1), 2389. DOI: 10.1038/s41467-023-38063-x.

(41) Galindo, G.; Maejima, D.; DeRoo, J.; Burlingham, S. R.; Fixen, G.; Morisaki, T.; Febvre, H. P.; Hasbrook, R.; Zhao, N.; Ghosh, S.;, et al. AI-assisted protein design to rapidly convert antibody sequences to intrabodies targeting diverse peptides and histone modifications. Sci Adv 2026, 12 (1), eadx8352. DOI: 10.1126/sciadv.adx8352.

(42) Gao, M.; Skolnick, J. Improved deep learning prediction of antigen-antibody interactions. Proc Natl Acad Sci U S A 2024, 121 (41), e2410529121. DOI: 10.1073/pnas.2410529121.

(43) Grassmann, G.; Di Rienzo, L.; Gosti, G.; Leonetti, M.; Ruocco, G.; Miotto, M.; Milanetti, E. Electrostatic complementarity at the interface drives transient protein-protein interactions. Sci Rep 2023, 13 (1), 10207. DOI: 10.1038/s41598-023-37130-z.

(44) Desantis, F.; Miotto, M.; Di Rienzo, L.; Milanetti, E.; Ruocco, G. Spatial organization of hydrophobic and charged residues affects protein thermal stability and binding affinity. Sci Rep 2022, 12 (1), 12087. DOI: 10.1038/s41598-022-16338-5.

(45) Hummer, A. M.; Abanades, B.; Deane, C. M. Advances in computational structure-based antibody design. Curr Opin Struct Biol 2022, 74, 102379. DOI: 10.1016/j.sbi.2022.102379.

(46) Baran, D.; Pszolla, M. G.; Lapidoth, G. D.; Norn, C.; Dym, O.; Unger, T.; Albeck, S.; Tyka, M. D.; Fleishman, S. J. Principles for computational design of binding antibodies. Proc Natl Acad Sci U S A 2017, 114 (41), 10900–10905. DOI: 10.1073/pnas.1707171114.

(47) Aguilar Rangel, M.; Bedwell, A.; Costanzi, E.; Taylor, R. J.; Russo, R.; Bernardes, G. J. L.; Ricagno, S.; Frydman, J.; Vendruscolo, M.; Sormanni, P. Fragment-based computational design of antibodies targeting structured epitopes. Sci Adv 2022, 8 (45), eabp9540. DOI: 10.1126/sciadv.abp9540.

(48) Bennett, N. R.; Watson, J. L.; Ragotte, R. J.; Borst, A. J.; See, D. L.; Weidle, C.; Biswas, R.; Yu, Y.; Shrock, E. L.; Ault, R.;, et al. Atomically accurate de novo design of antibodies with RFdiffusion. Nature 2026, 649 (8095), 183–193. DOI: 10.1038/s41586-025-09721-5.

(49) Bielska, W.; Jaszczyszyn, I.; Dudzic, P.; Janusz, B.; Chomicz, D.; Wrobel, S.; Greiff, V.; Feehan, R.; Adolf-Bryfogle, J.; Krawczyk, K. Applying computational protein design to therapeutic antibody discovery - current state and perspectives. Front Immunol 2025, 16, 1571371. DOI: 10.3389/fimmu.2025.1571371.

(50) Kingma, D. P.; Ba, J. Adam: a method for stochastic optimization. arXIV.org/abs/1412.6980. `2014`.

(51) Chrysostomou, C.; Seker, H. Novel protein weight matrix generated from amino acid indices. Annu Int Conf IEEE Eng Med Biol Soc 2015, 2015, 8181–8184. DOI: 10.1109/EMBC.2015.7320293.

(52) Gonnet, G. H.; Cohen, M. A.; Benner, S. A. Exhaustive matching of the entire protein sequence database. Science 1992, 256 (5062), 1443–1445. DOI: 10.1126/science.1604319.

(53) Jumper, J.; Evans, R.; Pritzel, A.; Green, T.; Figurnov, M.; Ronneberger, O.; Tunyasuvunakool, K.; Bates, R.; Zidek, A.; Potapenko, A.;, et al. Highly accurate protein structure prediction with AlphaFold. Nature 2021, 596 (7873), 583–589. DOI: 10.1038/s41586-021-03819-2.

(54) Lee, J. Y.; Lee, H. T.; Shin, W.; Chae, J.; Choi, J.; Kim, S. H.; Lim, H.; Won Heo, T.; Park, K. Y.; Lee, Y. J.;, et al. Structural basis of checkpoint blockade by monoclonal antibodies in cancer immunotherapy. Nat Commun 2016, 7, 13354. DOI: 10.1038/ncomms13354.

(55) Tan, S.; Zhang, H.; Chai, Y.; Song, H.; Tong, Z.; Wang, Q.; Qi, J.; Wong, G.; Zhu, X.; Liu, W. J.;, et al. An unexpected N-terminal loop in PD-1 dominates binding by nivolumab. Nat Commun 2017, 8, 14369. DOI: 10.1038/ncomms14369.

(56) Zak, K. M.; Kitel, R.; Przetocka, S.; Golik, P.; Guzik, K.; Musielak, B.; Domling, A.; Dubin, G.; Holak, T. A. Structure of the complex of human programmed death 1, PD-1, and its ligand PD-L1. Structure 2015, 23 (12), 2341–2348. DOI: 10.1016/j.str.2015.09.010.

(57) Schrödinger Maestro, S. R.-., LLC, New York, NY.

(58) Case, D. A.; Cheatham, T. E., 3rd; Darden, T.; Gohlke, H.; Luo, R.; Merz, K. M., Jr.; Onufriev, A.; Simmerling, C.; Wang, B.; Woods, R. J. The Amber biomolecular simulation programs. J Comput Chem 2005, 26 (16), 1668–1688. DOI: 10.1002/jcc.20290.

(59) Case, D. A.; Betz, R. M.; Botello-Smith, W.; Cerutti, D. S.; Cheatham, I. T. E.; Darden, T. A.; Duke, R. E.; Giese, T. J.; Gohlke, H.; Goetz, A. W.; et al. AMBER 2018. University of California, San Francisco 2018.

(60) Phillips, J. C.; Braun, R.; Wang, W.; Gumbart, J.; Tajkhorshid, E.; Villa, E.; Chipot, C.; Skeel, R. D.; Kale, L.; Schulten, K. Scalable molecular dynamics with NAMD. J Comput Chem 2005, 26 (16), 1781–1802. DOI: 10.1002/jcc.20289.

(61) Hopkins, C. W.; Le Grand, S.; Walker, R. C.; Roitberg, A. E. Long-time-step molecular dynamics through hydrogen mass repartitioning. Journal of Chemical Theory and Computation 2015, 11 (4), 1864–1874. DOI: 10.1021/ct5010406.

(62) Winn, M. D.; Ballard, C. C.; Cowtan, K. D.; Dodson, E. J.; Emsley, P.; Evans, P. R.; Keegan, R. M.; Krissinel, E. B.; Leslie, A. G.; McCoy, A.;, et al. Overview of the CCP4 suite and current developments. Acta Crystallogr D Biol Crystallogr 2011, 67 (Pt 4), 235–242. DOI: 10.1107/S0907444910045749.

(63) Humphrey, W.; Dalke, A.; Schulten, K. VMD: visual molecular dynamics. J Mol Graph 1996, 14 (1), 33–38, 27-38. DOI: 10.1016/0263-7855(96)00018-5.

(64) Wang, J.; Shi, Y.; Reiss, K.; Maschietto, F.; Lolis, E.; Konigsberg, W. H.; Lisi, G. P.; Batista, V. S. Structural insights into binding of remdesivir triphosphate within the replication-transcription complex of SARS-CoV-2. Biochemistry 2022, 61 (18), 1966–1973. DOI: 10.1021/acs.biochem.2c00341.

(65) Wang, J.; Shi, Y.; Reiss, K.; Allen, B.; Maschietto, F.; Lolis, E.; Konigsberg, W. H.; Lisi, G. P.; Batista, V. S. Insights into binding of single-stranded viral RNA template to the replication-transcription complex of SARS-CoV-2 for the priming reaction from molecular dynamics simulations. Biochemistry 2022, 61 (6), 424–432. DOI: 10.1021/acs.biochem.1c00755.

(66) Shi, Y.; Wang, J.; Batista, V. S. Translocation pause of remdesivir-containing primer/template RNA duplex within SARS-CoV-2’s RNA polymerase complexes. Front Mol Biosci 2022, 9, 999291. DOI: 10.3389/fmolb.2022.999291.

(67) Fredericks, A. M.; East, K. W.; Shi, Y.; Liu, J.; Maschietto, F.; Ayala, A.; Cioffi, W. G.; Cohen, M.; Fairbrother, W. G.; Lefort, C. T., et al. Identification and mechanistic basis of non-ACE2 blocking neutralizing antibodies from COVID-19 patients with deep RNA sequencing and molecular dynamics simulations. Front Mol Biosci 2022, 9, 1080964. DOI: 10.3389/fmolb.2022.1080964.

(68) Emsley, P.; Cowtan, K. Coot: model-building tools for molecular graphics. Acta Crystallogr D Biol Crystallogr 2004, 60 (Pt 12 Pt 1), 2126–2132. DOI: 10.1107/S0907444904019158.

(69) Adams, P. D.; Afonine, P. V.; Bunkoczi, G.; Chen, V. B.; Davis, I. W.; Echols, N.; Headd, J. J.; Hung, L. W.; Kapral, G. J.; Grosse-Kunstleve, R. W.;, et al. PHENIX: a comprehensive Python-based system for macromolecular structure solution. Acta Crystallogr D Biol Crystallogr 2010, 66 (Pt 2), 213–221. DOI: 10.1107/S0907444909052925.

(70) Delano, W. L. PyMol. Schrodinger, Inc. http://pymol.org/.

(71) Pollard, T. D. A guide to simple and informative binding assays. Mol Biol Cell 2010, 21 (23), 4061–4067. DOI: 10.1091/mbc.E10-08-0683.

(72) Du, Y. Binding curve viewer: visualizing the equilibrium and kinetics of protein-ligand binding and competitive binding. Journal of Chemical Information and Modeling 2024, 64 (10), 4180–4192. DOI: 10.1021/acs.jcim.4c00130.

(73) Chen, D.; Tan, S.; Zhang, H.; Wang, H.; He, W.; Shi, R.; Tong, Z.; Zhu, J.; Cheng, H.; Gao, S.;, et al. The FG Loop of PD-1 serves as a “hotspot" for therapeutic monoclonal antibodies in tumor immune checkpoint therapy. iScience 2019, 14, 113–124. DOI: 10.1016/j.isci.2019.03.017.

(74) Shrock, E. L.; Timms, R. T.; Kula, T.; Mena, E. L.; West, A. P., Jr.; Guo, R.; Lee, I. H.; Cohen, A. A.; McKay, L. G. A.; Bi, C.;, et al. Germline-encoded amino acid-binding motifs drive immunodominant public antibody responses. Science 2023, 380 (6640), eadc9498. DOI: 10.1126/science.adc9498.

(75) Wang, M.; Wang, J.; Wang, R.; Jiao, S.; Wang, S.; Zhang, J.; Zhang, M. Identification of a monoclonal antibody that targets PD-1 in a manner requiring PD-1 Asn58 glycosylation. Commun Biol 2019, 2, 392. DOI: 10.1038/s42003-019-0642-9.

(76) Lu, D.; Xu, Z. P.; Zhang, D.; Jiang, M.; Liu, K. F.; He, J. H.; Ma, D. L.; Ma, X. P.; Tan, S. G.; Gao, G. F.;, et al. PD-1 N58-Glycosylation-Dependent Binding of Monoclonal Antibody Cemiplimab for Immune Checkpoint Therapy. Front Immunol 2022, 13. DOI: ARTN 826045 10.3389/fimmu.2022.826045.

(77) Liu, K.; Tan, S.; Jin, W.; Guan, J.; Wang, Q.; Sun, H.; Qi, J.; Yan, J.; Chai, Y.; Wang, Z.;, et al. N-glycosylation of PD-1 promotes binding of camrelizumab. EMBO Rep 2020, 21 (12), e51444. DOI: 10.15252/embr.202051444.

(78) Liu, H.; Guo, L.; Zhang, J.; Zhou, Y.; Zhou, J.; Yao, J.; Wu, H.; Yao, S.; Chen, B.; Chai, Y.;, et al. Glycosylation-independent binding of monoclonal antibody toripalimab to FG loop of PD-1 for tumor immune checkpoint therapy. MAbs 2019, 11 (4), 681–690. DOI: 10.1080/19420862.2019.1596513.

(79) Wang, J.; Skeens, E.; Arantes, P. R.; Maschietto, F.; Allen, B.; Kyro, G. W.; Lisi, G. P.; Palermo, G.; Batista, V. S. Structural basis for reduced dynamics of three engineered HNH endonuclease Lys-to-Ala mutants of the Cas9 enzyme. Biochemistry 2022, 61, 785–794.

(80) Pennington, L. F.; Tarchevskaya, S.; Brigger, D.; Sathiyamoorthy, K.; Graham, M. T.; Nadeau, K. C.; Eggel, A.; Jardetzky, T. S. Structural basis of omalizumab therapy and omalizumab-mediated IgE exchange. Nat Commun 2016, 7, 11610. DOI: 10.1038/ncomms11610.

(81) Lieu, R.; Antonysamy, S.; Druzina, Z.; Ho, C.; Kang, N. R.; Pustilnik, A.; Wang, J.; Atwell, S. Rapid and robust antibody Fab fragment crystallization utilizing edge-to-edge beta-sheet packing. PLoS One 2020, 15 (9), e0232311. DOI: 10.1371/journal.pone.0232311.

(82) Koide, S.; Sidhu, S. S. The importance of being tyrosine: lessons in molecular recognition from minimalist synthetic binding proteins. ACS Chem Biol 2009, 4 (5), 325–334. DOI: 10.1021/cb800314v.

(83) Fellouse, F. A.; Wiesmann, C.; Sidhu, S. S. Synthetic antibodies from a four-amino-acid code: a dominant role for tyrosine in antigen recognition. Proc Natl Acad Sci U S A 2004, 101 (34), 12467–12472. DOI: 10.1073/pnas.0401786101.

(84) Birtalan, S.; Zhang, Y.; Fellouse, F. A.; Shao, L.; Schaefer, G.; Sidhu, S. S. The intrinsic contributions of tyrosine, serine, glycine and arginine to the affinity and specificity of antibodies. J Mol Biol 2008, 377 (5), 1518–1528. DOI: 10.1016/j.jmb.2008.01.093.

(85) Chen, C.; Roberts, V. A.; Rittenberg, M. B. Generation and analysis of random point mutations in an antibody CDR2 sequence - Many mutated antibodies lose their ability to bind antigen. Journal of Experimental Medicine 1992, 176 (3), 855–866. DOI: DOI 10.1084/jem.176.3.855.

(86) Wiehe, K.; Bradley, T.; Meyerhoff, R. R.; Hart, C.; Williams, W. B.; Easterhoff, D.; Faison, W. J.; Kepler, T. B.; Saunders, K. O.; Alam, S. M.;, et al. Functional relevance of improbable antibody mutations for HIV broadly neutralizing antibody development. Cell Host Microbe 2018, 23 (6), 759–765 e756. DOI: 10.1016/j.chom.2018.04.018.

(87) Oyama, H.; Kiguchi, Y.; Morita, I.; Yamamoto, C.; Higashi, Y.; Taguchi, M.; Tagawa, T.; Enami, Y.; Takamine, Y.; Hasegawa, H.;, et al. Seeking high-priority mutations enabling successful antibody-breeding: systematic analysis of a mutant that gained over 100-fold enhanced affinity. Sci Rep 2020, 10 (1), 4807. DOI: 10.1038/s41598-020-61529-7.

(88) Fernandez-Quintero, M. L.; Kokot, J.; Waibl, F.; Fischer, A. M.; Quoika, P. K.; Deane, C. M.; Liedl, K. R. Challenges in antibody structure prediction. MAbs 2023, 15 (1), 2175319. DOI: 10.1080/19420862.2023.2175319.

(89) AlQuraishi, M. Machine learning in protein structure prediction. Curr Opin Chem Biol 2021, 65, 1–8. DOI: 10.1016/j.cbpa.2021.04.005.

(90) Abanades, B.; Georges, G.; Bujotzek, A.; Deane, C. M. ABlooper: fast accurate antibody CDR loop structure prediction with accuracy estimation. Bioinformatics 2022, 38 (7), 1877–1880. DOI: 10.1093/bioinformatics/btac016.

(91) Maier, J. K.; Labute, P. Assessment of fully automated antibody homology modeling protocols in molecular operating environment. Proteins 2014, 82 (8), 1599–1610. DOI: 10.1002/prot.24576.

(92) Zhu, K.; Day, T.; Warshaviak, D.; Murrett, C.; Friesner, R.; Pearlman, D. Antibody structure determination using a combination of homology modeling, energy-based refinement, and loop prediction. Proteins 2014, 82 (8), 1646–1655. DOI: 10.1002/prot.24551.

(93) Ruffolo, J. A.; Sulam, J.; Gray, J. J. Antibody structure prediction using interpretable deep learning. Patterns (N Y) 2022, 3 (2), 100406. DOI: 10.1016/j.patter.2021.100406.

(94) Abanades, B.; Wong, W. K.; Boyles, F.; Georges, G.; Bujotzek, A.; Deane, C. M. ImmuneBuilder: Deep-Learning models for predicting the structures of immune proteins. Commun Biol 2023, 6 (1), 575. DOI: 10.1038/s42003-023-04927-7.

(95) Baek, M.; DiMaio, F.; Anishchenko, I.; Dauparas, J.; Ovchinnikov, S.; Lee, G. R.; Wang, J.; Cong, Q.; Kinch, L. N.; Schaeffer, R. D.;, et al. Accurate prediction of protein structures and interactions using a three-track neural network. Science 2021, 373 (6557), 871–876. DOI: 10.1126/science.abj8754.

(96) Gainza, P.; Wehrle, S.; Van Hall-Beauvais, A.; Marchand, A.; Scheck, A.; Harteveld, Z.; Buckley, S.; Ni, D.; Tan, S.; Sverrisson, F.;, et al. De novo design of protein interactions with learned surface fingerprints. Nature 2023, 617 (7959), 176–184. DOI: 10.1038/s41586-023-05993-x.

(97) Wang, J.; Maschietto, F.; Qiu, T.; Arantes, P. R.; Skeens, E.; Palermo, G.; Lisi, G. P.; Batista, V. S. Substrate-independent activation pathways of the CRISPR-Cas9 HNH nuclease. Biophys J 2023, 122 (24), 4635–4644. DOI: 10.1016/j.bpj.2023.11.005.

(98) Lin, D. Y.; Tanaka, Y.; Iwasaki, M.; Gittis, A. G.; Su, H. P.; Mikami, B.; Okazaki, T.; Honjo, T.; Minato, N.; Garboczi, D. N. The PD-1/PD-L1 complex resembles the antigen-binding Fv domains of antibodies and T cell receptors. Proc Natl Acad Sci U S A 2008, 105 (8), 3011–3016. DOI: 10.1073/pnas.0712278105.

(99) Lazar-Molnar, E.; Yan, Q.; Cao, E.; Ramagopal, U.; Nathenson, S. G.; Almo, S. C. Crystal structure of the complex between programmed death-1 (PD-1) and its ligand PD-L2. Proc Natl Acad Sci U S A 2008, 105 (30), 10483–10488. DOI: 10.1073/pnas.0804453105 From NLM Medline.

(100) Cheng, X.; Veverka, V.; Radhakrishnan, A.; Waters, L. C.; Muskett, F. W.; Morgan, S. H.; Huo, J.; Yu, C.; Evans, E. J.; Leslie, A. J.;, et al. Structure and interactions of the human programmed cell death 1 receptor. J Biol Chem 2013, 288 (17), 11771–11785. DOI: 10.1074/jbc.M112.448126.

(101) Tang, S. G.; Kim, P. S. A high-affinity human PD-1/PD-L2 complex informs avenues for small-molecule immune checkpoint drug discovery. P Natl Acad Sci USA 2019, 116 (49), 24500–24506. DOI: 10.1073/pnas.1916916116.

(102) Na, Z.; Yeo, S. P.; Bharath, S. R.; Bowler, M. W.; Balikci, E.; Wang, C. I.; Song, H. Structural basis for blocking PD-1-mediated immune suppression by therapeutic antibody pembrolizumab. Cell Res 2017, 27 (1), 147–150. DOI: 10.1038/cr.2016.77.

(103) Paulk, A. M.; Williams, R. L.; Liu, C. C. Rapidly Inducible Yeast Surface Display for Antibody Evolution with OrthoRep. ACS Synth Biol 2024, 13 (8), 2629–2634. DOI: 10.1021/acssynbio.4c00370.

(104) Zhang, L.; Ding, Y.; Wang, Q.; Pan, W.; Wei, Z.; Smith, P. A.; Yang, X. Preclinical immunological characterization of rademikibart (CBP-201), a next-generation human monoclonal antibody targeting IL-4Ralpha, for the treatment of Th2 inflammatory diseases. Sci Rep 2023, 13 (1), 12411. DOI: 10.1038/s41598-023-39311-2.

